# The effect of dipeptide repeat proteins on FUS/TDP43-RNA condensation in C9orf72 ALS/FTD

**DOI:** 10.1101/2024.05.21.595197

**Authors:** Mark D. Driver, Jasper Postema, Patrick R. Onck

**Affiliations:** Zernike Institute for Advanced Materials, University of Groningen, Groningen, The Netherlands

**Keywords:** phase separation, ALS, C9orf72, DPR, coarse-grained molecular dynamics

## Abstract

Condensation of RNA binding proteins (RBPs) with RNA is essential for cellular function. The most common familial cause of the diseases ALS and FTD are C9orf72 repeat expansion disorders that produce dipeptide repeat proteins (DPRs). We explore the hypothesis that DPRs disrupt the native condensation behaviour of RBPs and RNA through molecular interactions resulting in toxicity. FUS and TDP43 are two RBPs known to be affected in ALS/FTD. We use our previously developed 1-bead-per-amino acid and a newly developed 3-bead-per-nucleic acid molecular dynamics model to explore ternary phase diagrams of FUS/TDP43-RNA-DPR systems. We show that the most toxic arginine containing DPRs (R-DPRs), can disrupt the RBP condensates through cation-*π* interactions, and can strongly sequester RNA through electrostatic interactions. The native droplet morphologies are already modified at small additions of R-DPRs leading to non-native FUS/TDP43-encapsulated condensates with a marbled RNA/DPR core.

## INTRODUCTION

Amyotrophic lateral sclerosis (ALS) and frontotemporal dementia (FTD) are striking neurodegenerative diseases that have distinct clinical presentations. ALS predominantly affects motor neurones, leading to decline in muscular control and function typically leading to death within 2-3 years after the onset of symptoms (1–5). On the other hand FTD affects the frontal and temporal lobes of the brain leading to a decline in cognitive function and memory, with a similarly short lifespan (6–8). Exploration of genomic data has led to the identification of multiple possible causes for ALS (2, 9) and FTD (5, 7, 8). Intriguingly, the most frequent cause of familial ALS, the G4C2 hexanucleotide repeat expansion in C9orf72, is also a common cause of FTD (5, 10–14). Typically up to 20 repeats of G4C2 are present in healthy individuals, whereas up to hundreds or thousands of repeats can be present in patients with C9orf72-mediated ALS/FTD (C9-ALS/FTD) (4). Translation of the RNA transcripts of this repeat expansion can produce five different dipeptide repeat proteins (DPRs): poly-PR, poly-GR, poly-GA, poly-GP, and poly-PA (15–18). The onset of toxicity from C9-ALS/FTD can be caused by a loss of function mechanism by the deactivation of the C9orf72 protein by the mutation (11) or by differential C9orf72 protein transcription rates affecting protein efficacy (19). Alternatively, the toxicity could arise from the gain of function caused by either direct interactions of the RNA transcripts produced (1, 20–23) or by the interactions of the DPRs produced by translation of the RNA transcripts (15, 16, 24, 25).

The highest level of toxicity is exhibited by the arginine-rich DPRs polyPR and polyGR (R-DPRs) which is shown in both animal and cell models (9, 26–29) with greater toxicity seen with increased repeat lengths in cells (30). A wide variety of cellular defects is hypothesised to be the result of R-DPRs such as compromised nuclear transport (31–33). Recent evidence also links the production of R-DPRs with the disruption of membraneless organelle (MLO) function (22, 24, 27, 31). MLOs, also called biological condensates, are formed by (liquid-liquid) phase separation (34–42) driven by protein-protein and protein-RNA interactions, often through intrinsically disordered regions (IDRs) of proteins (43–47). Indeed, it has previously been shown that R-DPRs interact with RBPs (24), including disruption of the LLPS of FUS (48–51). Interestingly, the positive charge of the R-DPRs can also lead to their direct interaction with RNA to form condensates (32, 47).

Of the DPRs which do not contain arginine (polyGA, polyPA, polyGP), the ΔR-DPRs, only polyGA has been found to show toxicity (30, 32, 52, 53). In fact, polyPA has been found to ameliorate polyGA toxicity when coexpressed (52, 53). The ΔR-DPRs do not exhibit the same ability to interact with IDRs that are common in RBPs (24), indicating they are less able to disrupt RBP function through the modification of LLPS. The effect of polyGA on cleavage rates of TDP43 (an RBP) indicates a mechanism that is potentially unrelated to MLO formation (52, 53).

In this work we explore the C9-ALS/FTD toxicity involving the disruption of RBP MLO function by DPRs. It can be hypothesised that R-DPR disruption occurs via two possible pathways. The first is related to the direct interaction of R-DPRs with RBPs to disrupt native RBP LLPS. The second is related to an indirect mechanism in which R-DPR interactions with RNA disrupt the native formation of RBP-RNA MLOs. The ΔR-DPRs, on the other hand, are hypothesised to not interact with RNA, and to have only a minimal effect on RBP condensates. To test these hypotheses, we use coarse grained molecular dynamics (CGMD) to explore the effect of DPRs on the LLPS of FUS and TDP43 in the presence and absence of RNA.

The paper is organised as follows. First we carry out a detailed examination of the ternary system of FUS with U_40_ (a simple RNA homopolymer) and PR_60_ (an R-DPR above the repeat length toxicity threshold). Next, the effect of R-DPR sequence composition and length on FUS-U_40_ interactions are discussed. Then we compare FUS with TDP43 by carrying out simulations on the ternary TDP43, U_40_, DPR system and we close off by studying the interactions of ΔR-DPRs on both FUS and TDP43. Our results show that R-DPRs co-condensate with FUS and TDP43 forming marbled droplets. More strikingly is the effect on native RBP-RNA condensation with R-DPRs strongly disrupting FUS/RNA and TDP43/RNA droplet morphologies, already at small additions of R-DPRs, modulating condensate architectures from pure FUS or TDP43 into marbled RNA/R-DPR domains infiltrating the RBP droplets.

## RESULTS

### Aromatic and cation-*π* interactions drive FUS LLPS

Fig. 1 shows that pure FUS (residues 1-267) undergoes LLPS, whereas pure U_40_ and pure PR_60_ do not. The driving forces for the LLPS of FUS observed in Fig. 1 can be understood by considering the intermolecular interactions (shown in Fig. 2). FUS (residues 1-267) contains two domains: residues 1-167 are the PLD (Prion-like domain) which is QGSY-rich and residues 168-267 are the RGG1 domain (which contains RGG repeat motifs). Fig. 2A shows that FUS undergoes LLPS through the interaction of aromatic tyrosine residues in the PLD domain that interact with other tyrosine residues in the PLD domain as well as through cation-*π* interactions with the arginine residues of the RGG1 domain. Hydrophilic residues in FUS (glycine, serine, threonine, and glutamine) show a large number of contacts in Fig. 2B, due to their high abundances in the PLD and RGG1 domains, yet have a significantly lower number of per residue contacts than tyrosine or arginine. Fig. 2C shows that the most important sticker interactions in FUS LLPS are the aromatic interactions, with a significant contribution from the cation-*π* interactions between the two FUS domains. This is in agreement with *in vitro* studies that indicated FUS LLPS to be driven by the PLD (54, 55).

**Figure 1:**
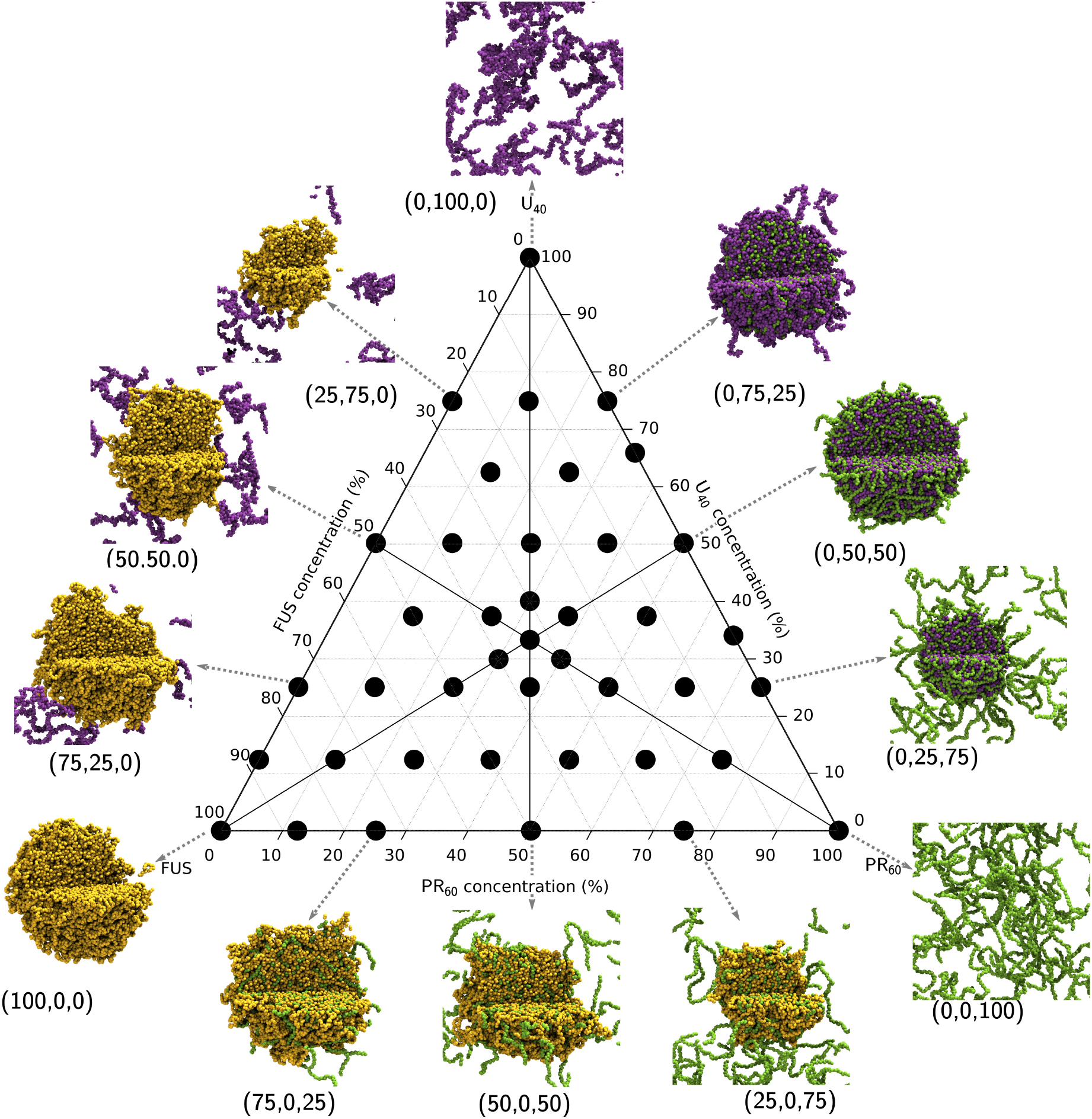
Ternary phase diagram showing the simulations undertaken in this work. Simulations were all run with an amino acid concentration of 80000 *μM*. The total composition of a system is defined relative to 120 FUS molecules, 267 U_40_ molecules, and 267 polyPR_60_ molecules, to give percentage compositions described in the image. The end frame of 3*μ*s of simulation is displayed as a representative state. FUS molecules are coloured in yellow, U_40_ molecules are coloured in purple, and 267 polyPR_60_ molecules are coloured in green. Composition as a percentage is shown next to the droplets in brackets: (% FUS, % U_40_, % polyPR_60_).

**Figure 2:**
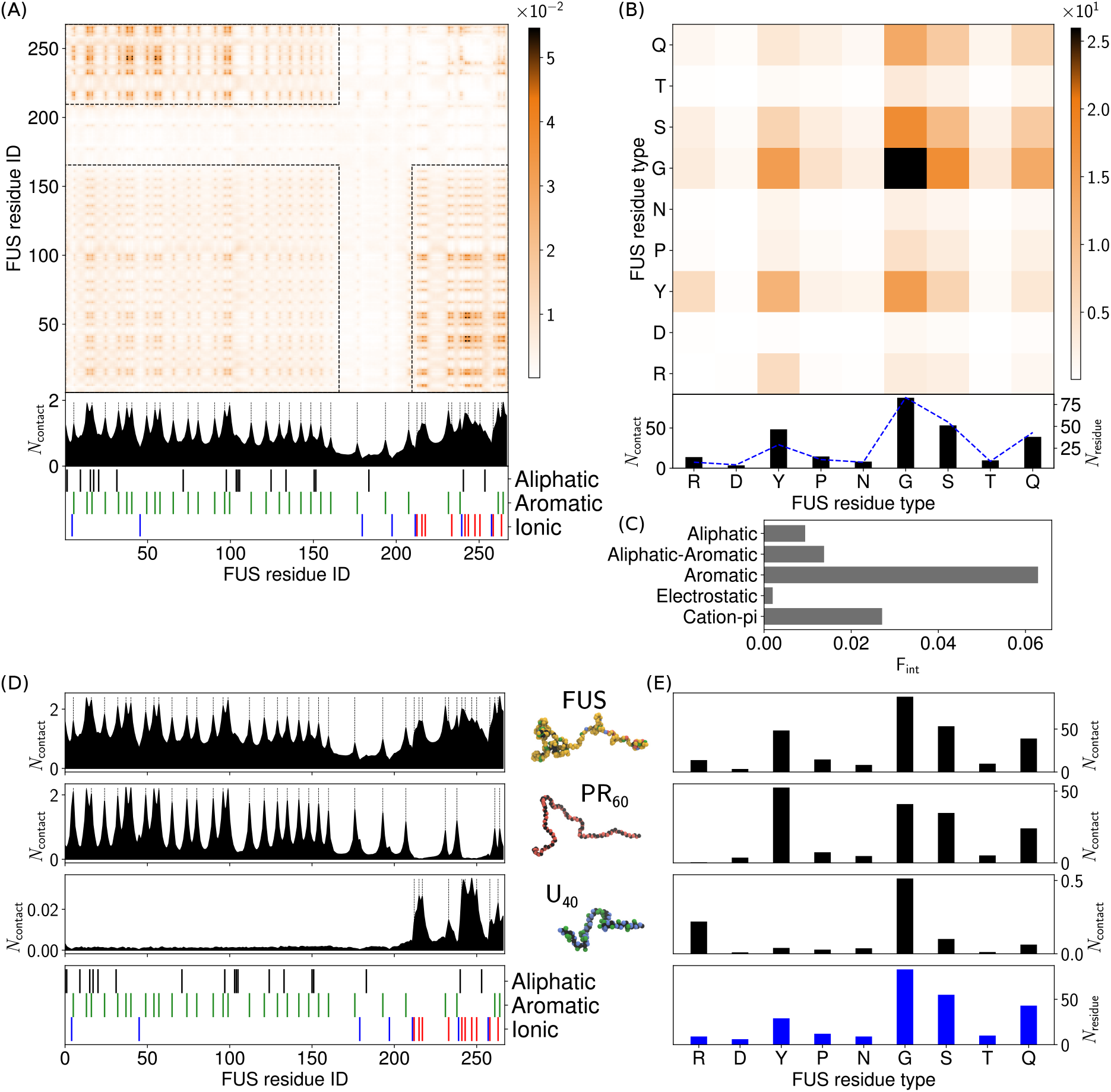
Intermolecular contact information for FUS interactions. (A) FUS intermolecular contact map by residue index for 100% FUS at 150 mM ion concentration and 300 K. The contacts are averaged in time and normalised by the number of molecules in the simulation (see section 3.2 in the SI for more details). A 1D contact profile (summation of the 2D map) is included below the contact map to show the total interactions per residue index (*N*_contact_). The black dashed lines highlight the location of tyrosine and arginine residues (which correspond to the peaks in the 1D profile). (B) Intermolecular contact map by residue type for FUS. The contact map in (B) is similar to the contact map by residue index in (A), but aggregated by residue type. A 1D contact profile (summation of the 2D map), are also included below (*N*_contact_) together with the abundance for the residues (*N*_residue_) shown by blue dashed lines. (C) Interaction summary for a droplet simulation with 100% FUS at 150 mM and 300 K. The fraction of interactions, F_int_, are aggregated by type and normalised by the total number of interactions. Aromatic and aliphatic interactions denote aromatic-aromatic and aliphatic-aliphatic interactions respectively. This convention is used throughout this work. Details of the contact definitions can be found in section 3.2 of the SI. (D) Intermolecular contact summaries by FUS residue index (bottom) for FUS with FUS (first row), PR_60_ (second row) and U_40_ (third row) (E) Intermolecular contact summaries by FUS residue type (bottom) for FUS with FUS (first row), PR_60_ (second row) and U_40_ (third row).

### FUS interacts with PR_60_ through tyrosines in the PLD and with U_40_ by arginines in the RGG domain

Fig. 1 shows that upon addition of U_40_ to FUS, FUS condensates are still observed with interactions between the FUS and U_40_ only observed with the condensate surface, and no U_40_ is observed to enter the core of the droplets. This is in contrast to the behaviour of FUS with PR_60_, which show FUS droplets with PR_60_ inclusions that result in a marbled pattern in the droplets, and excess PR_60_ in the dilute phase. The heterotypic interactions of FUS with PR_60_ are dominated by cation-*π* interactions of the tyrosine residues in FUS interacting with the arginine residues in PR_60_ (see 1D contact summaries in Fig. 2D, with full contact maps in Figs. S5-S14 in the SI). These cation-*π* contacts are concentrated in the FUS PLD domain, which has the majority of the tyrosine residues in FUS. Fig. 2D shows that the observed number of contacts of the tyrosines in the RGG1 domain is lower than the contacts of the tyrosines in the PLD domain.

FUS-U_40_ contacts however, predominantly originate from the RGG1 domain of FUS, driven by electrostatic interactions of the FUS arginine residues and the phosphate groups in U_40_ (Figs. 2D and 2E). Significant numbers of U_40_ contacts with glycine are also observed, due to high abundance of glycine in the RGG domain (glycine accounts for 50% of the residues in the RGG1 domain) and the proximity of glycine to arginine in the RGG motifs that drive RNA binding. Interactions of the FUS PLD with U_40_ are significantly weaker due to the hydrophilicity of the phosphate group in U_40_ resulting in repulsive interactions with the tyrosines and polar residues in the PLD. The observed behaviour of U_40_ being unable to enter the FUS condensates in the binary FUS-U_40_ mixtures can be explained by these different FUS domain interactions, and it instead only interacts with exposed FUS RGG1 on the surface of the droplets. This behaviour is in agreement with *in vitro* results that have shown that FUS RGG domain interactions with RNA can promote LLPS (56, 57).

### Electrostatic interactions drive complex coacervation of U_40_ with PR_60_

The mixing of U_40_ and PR_60_ produces marbled condensates and strong LLPS, with mixtures of (0,50,50) and (0,67,33) forming only a single droplet with no dilute phase (see images in Fig. 1). A dilute phase of U_40_ exists at higher U_40_ proportions (see (0,75,25) in Fig. 1), whereas a dilute phase of PR_60_ exists at higher PR_60_ fractions (see (0,25,75) in Fig. 1). U_40_ and PR_60_ are polyelectrolyte polymers which are negatively or positively charged respectively, resulting in repulsive homotypic interactions and thus are unable to undergo simple coacervation. Instead, complex coacervation is observed between U_40_ and PR_60_, (shown in Fig. 1). The sequence uniformity of U_40_ and PR_60_ result in contact maps with a large degree of homogeneity (see Figs. S13 and S14 in the SI), with electrostatic interactions between arginine of PR_60_ and the phosphate group of U_40_ dominating the interactions. Interestingly, a large number of interactions between the proline of PR_60_ and the uracil bead of U_40_ are also observed with fewer contacts with the ribose group, indicating that the PR_60_ strands wrap around the RNA such that proline-uracil contacts are formed instead of proline-ribose interactions. Based on the hydrophobicity of the ribose (*ϵ*_*i*_=0.755) and uracil (*ϵ*_*i*_=0.382) groups, proline (*ϵ*_*i*_=0.67) interactions with ribose are energetically preferred, indicating a relatively repulsive driving force for the observed proline-uracil interactions.

### Competitive heterotypic interactions drive the formation of complex morphologies in ternary FUS-U_40_-PR_60_ systems

Interestingly, when we simulate compositions with all three components (FUS, U_40_, and PR_60_) we observed a shift in droplet morphology related to the system composition. Four distinct regions of the phase diagram exist in Fig. 3: no LLPS observed (PR_60_ and U_40_ corners), full LLPS (white) where no significant dilute phase is present, U_40_ in the dilute phase (purple), and PR_60_ in the dilute phase (green). It is important to note that the intermolecular contact maps for homotypic and heterotypic interactions of FUS, U_40_ and PR_60_ all display the same interaction patterns, irrespective of the composition (see section 6 in the SI), with only the relative number of interactions changing.

**Figure 3:**
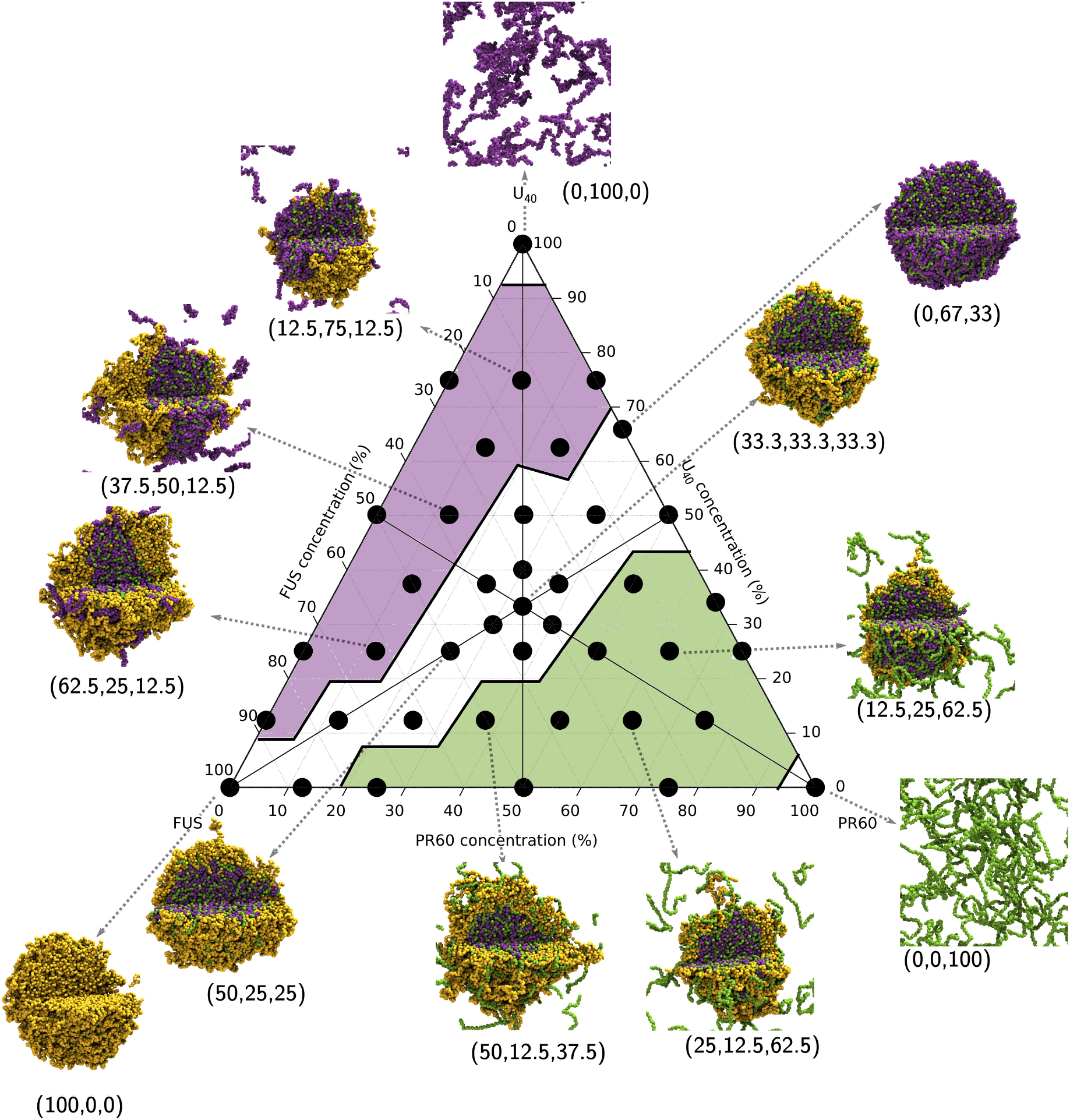
Ternary phase diagram showing the simulations undertaken in this work. Simulations were all run with an amino acid concentration of 80000 *μM*. The total composition of a system is defined relative to 120 FUS molecules, 267 U_40_ molecules, and 267 polyPR_60_ molecules, to give percentage compositions described in the image. The end frame of 3*μ*s of simulation is displayed as a representative state. FUS molecules are coloured in yellow, U_40_ molecules are coloured in purple, and 267 polyPR_60_ molecules are coloured in green. Composition as a percentage is shown next to the droplets in brackets: (% FUS, % U_40_, % polyPR_60_). The phase diagram is divided into four regions: no LLPS observed (PR_60_ and U_40_ corners), full LLPS (white) where no significant dilute phase is present, U_40_ in the dilute phase (purple), and PR_60_ in the dilute phase (green).

The binary mixture of U_40_ and PR_60_ at (0, 67, 33) on the right hand edge of the white region in Fig. 3 is charge neutral and all molecules are in the condensed phase. A lower amount of PR_60_ is visible in the (0, 67, 33) droplet than in (0, 50, 50) droplet where there is a net positive charge across the biomolecules, and an equal number of PR_60_ and U_40_ particles in the simulation. As we traverse across the triangle towards the FUS corner at (100, 0, 0), we see that the FUS forms a coating on the surface of the U_40_-PR_60_ condensates that eventually completely covers the U_40_-PR_60_ droplet surface at (33.3, 33.3, 33.3), before the FUS layer thickens and the U_40_-PR_60_ core disappears at (100, 0, 0).

In the green (lower right) region of the phase diagram in Fig. 3, we observed that the droplets have a similar structure to those in the white region, with the notable difference that the increased PR_60_ fraction results in the outer shell containing a mixture of FUS and PR_60_. The increased PR_60_ concentration also leads to the presence of a dilute phase of PR_60_ which are not incorporated into the condensate. This is due to the electrostatic intermolecular repulsion of the PR_60_ molecules. In the purple (upper left) region of the phase diagram in Fig. 3, higher U_40_ concentrations result in the presence of U_40_ in the dilute phase. The presence of U_40_, which is not soluble in the FUS PLD (56) leads to a situation where FUS no longer completely coats the U_40_-PR_60_ condensate at higher FUS concentrations. Instead, FUS forms a condensate that is clustered, to enable U_40_ exchange between the dilute and condensed phases.

In the binary compositions of FUS-U_40_ in Fig. 1 where U_40_ interacts with the FUS condensate surface, the addition of PR_60_ has led to a change in the morphology of the condensate droplets. The resultant ternary mixtures show FUS forming a shell around U_40_-PR_60_ droplets. Within a core-shell architecture the shell condensate has a larger surface area to volume ration than the core condensate. With the increasing evidence that interfaces are an important site for aggregate initiation, the PR_60_ induced morphological change could provide a potential mechanism for R-DPR toxicity, where the morphological changes reduce the barriers to aggregation by increasing the reactive surface area.

All in all, our results show that already at small additions of PR_60_, strong perturbations of droplet morphology are observed, with marbled U_40_-PR_60_ domains infiltrating the FUS condensates (purple region of Fig. 3). At larger PR_60_ additions, the U_40_-PR_60_ domains are fully engulfed by FUS, forming a FUS coated U_40_-PR_60_ marbled core (white region of Fig. 3). Finally, at high PR_60_ concentrations no strong droplet perturbations are observed with excess PR_60_ resulting in a dilute phase.

### R-DPR length influences the observed condensate morphologies

Fig. 3 for the FUS-U_40_-PR_60_ system provides a detailed morphological reference case for the investigation of R-DPR sequence modifications on condensate behaviour. To explore the effect of sequence length and sequence composition, simulations using PR_30_, PR_15_, GR_60_, GR_30_, and GR_15_ were completed. The interactions of all R-DPRs with FUS and RNA are driven by arginine residues in the R-DPR, with cation-*π* interactions formed with FUS and electrostatic interactions formed with U_40_ (see Contact maps in Figs. S5-S14 in the SI).

The same colouring scheme as Fig. 3 for the annotated triangles of PR_60_, PR_30_, PR_15_, GR_60_, GR_30_, and GR_15_ was used to summarise the results in Fig. S2. Fig. S2 shows that there was no observable difference in condensate behaviour for R-DPRs of the same length (PR or GR). As the number of repeat units was decreased a reduction in the size of the central full LLPS (white) region was observed. This can be understood by two complementary factors: 1) the reduction in R-DPR multivalency, and 2) increased inter-R-DPR repulsion. The shorter R-DPR molecules are forming fewer interactions with FUS or U_40_, reducing their ability to glue FUS and U_40_ together. An increase in R-DPR molecules is required to maintain the same number R-DPR residues in simulations with the same R-DPR mass fraction. Without the additional covalent bonds holding these smaller fragments together, the electrostatic repulsion causes increased R-DPR repulsive interactions that lowers the stability of large U_40_-R-DPR droplets.

### Aliphatic residues drive TDP43 LLPS with aromatic residues enabling co-condensation with R-DPRs

TDP43 is another RBP that undergoes LLPS driven by the C terminal IDR (58–61). The TDP43 IDR contains many similar sequence features to the FUS PLD. Like FUS, the TDP43 (IDR, omitted further on for clarity) contains several aromatic residues (phenylalanine and tryptophan, rather than tyrosine), and has a low number of ionic residues (6/140 with a net charge of +2). TDP43 also contains a higher fraction of aliphatic residues. Fig. 4 shows the intermolecular contact maps for TDP43 interactions. Examination of the TDP43-TDP43 intermolecular contacts in Fig. 4A shows peaks in the contacts around aromatic residues. The 1D interaction summary (Fig. 4A) shows a broad area of interactions between residues 40 and 67, corresponding to a tract high in aliphatic residues. The aliphatic residues alanine, leucine, methionine, and proline in Fig. 4B show a significant number of contacts, that make aliphatic interactions the most important driving force for TDP43 intermolecular interactions (Fig. 4C), compared to aromatic contacts in FUS (2). The hydrophilic residues (glycine, serine, asparagine, and glutamine), which account for 63% of the residues in TDP43) show a large number of contacts in Fig. 4B, due to their high abundances in the CTD domain, yet have a significantly lower number of per residue contacts like the case of FUS.

**Figure 4:**
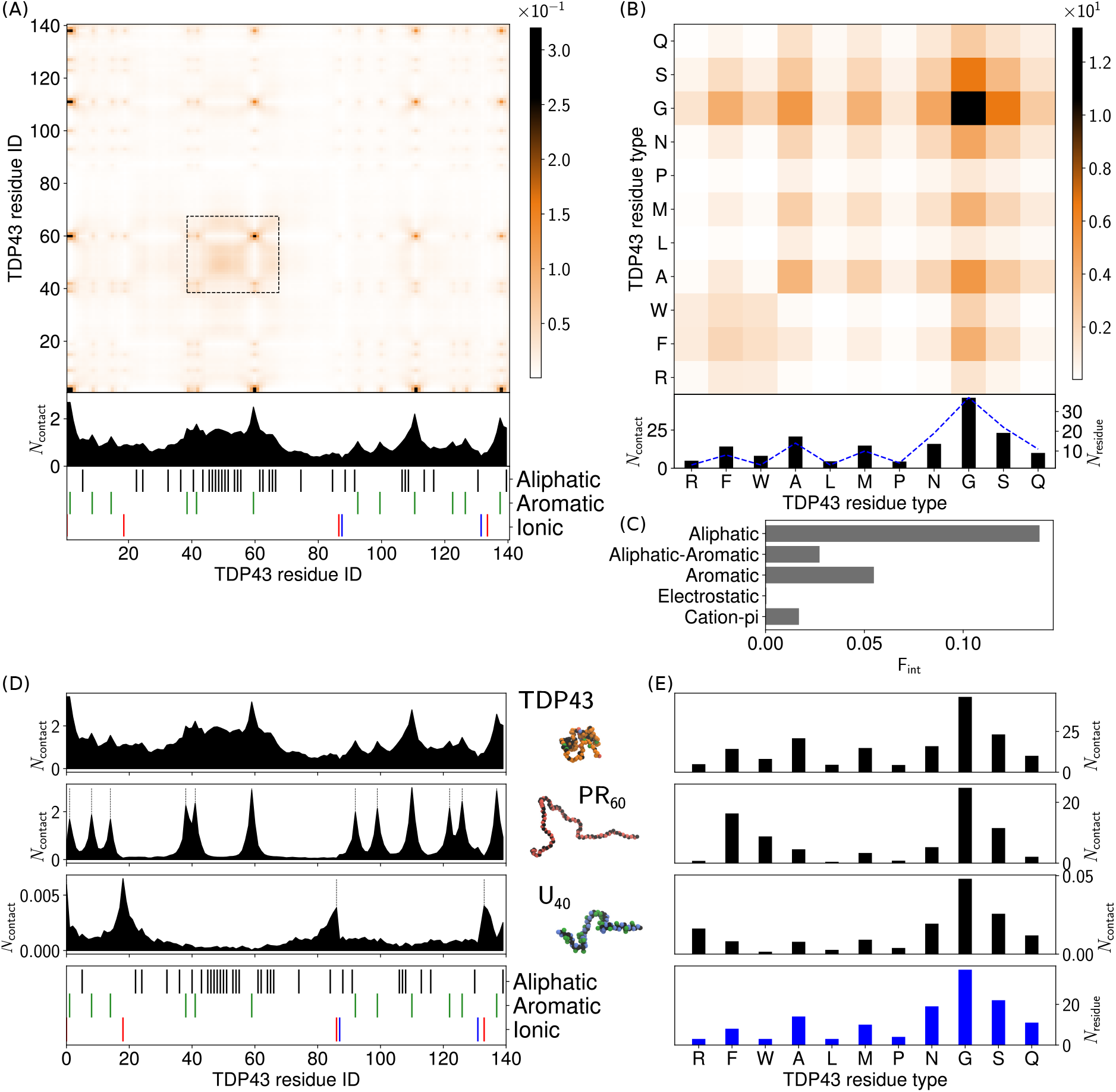
Intermolecular contact maps for TDP43 interactions in a single-component droplet. (A) TDP43 intermolecular contact map by residue index for 100% TDP43 at 150 mM ion concentration and 300 K. The contacts are averaged in time and normalised by the number of molecules in the simulation (see section 3.2 in the SI for more details). A 1D contact profile (summation of the 2D map) is included below the contact map to show the total interactions per residue index (*N*_contact_). (B) Intermolecular contact map by residue type for TDP43. The contact map in (B) is similar to the contact map by residue index in (A), but aggregated by residue type. A 1D contact profile (summation of the 2D map), are also included below (*N*_contact_) together with the abundance for the residues (*N*_residue_) shown by blue dashed lines. (C) Interaction summary for a droplet simulation with 100% TDP43 at 150 mM and 300 K. The fraction of interactions, F_int_, are aggregated by type and normalised by the total number of interactions. Aromatic and aliphatic interactions denote aromatic-aromatic and aliphatic-aliphatic interactions respectively. This convention is used throughout this work. Details of the contact definitions can be found in section 3.2 of the SI. (D) Intermolecular contact summaries by TDP43 residue index (bottom) for TDP43 with TDP43 (first row), PR_60_ (second row) and U_40_ (third row) (E) Intermolecular contact summaries by TDP43 residue type (bottom) for TDP43 with TDP43 (first row), PR_60_ (second row) and U_40_ (third row).

If we now consider the ability of the TDP43 to interact with U_40_ and PR_60_, we see similar behaviour to the FUS PLD. In Fig. 4D and E we see that PR_60_ interactions are concentrated at the aromatic residues in TDP43, resulting in PR_60_ inclusions in TDP43 condensates (Fig. 5). As expected, minimal interactions are seen with U_40_ (Fig. 4D and E), with the 1D summary showing aggregate contacts two orders of magnitude lower than TDP43 with TDP43 or PR_60_. This results in TDP43 droplets excluding U_40_. Therefore without the RRM domains RNA binding of TDP43 is significantly curtailed (60).

**Figure 5:**
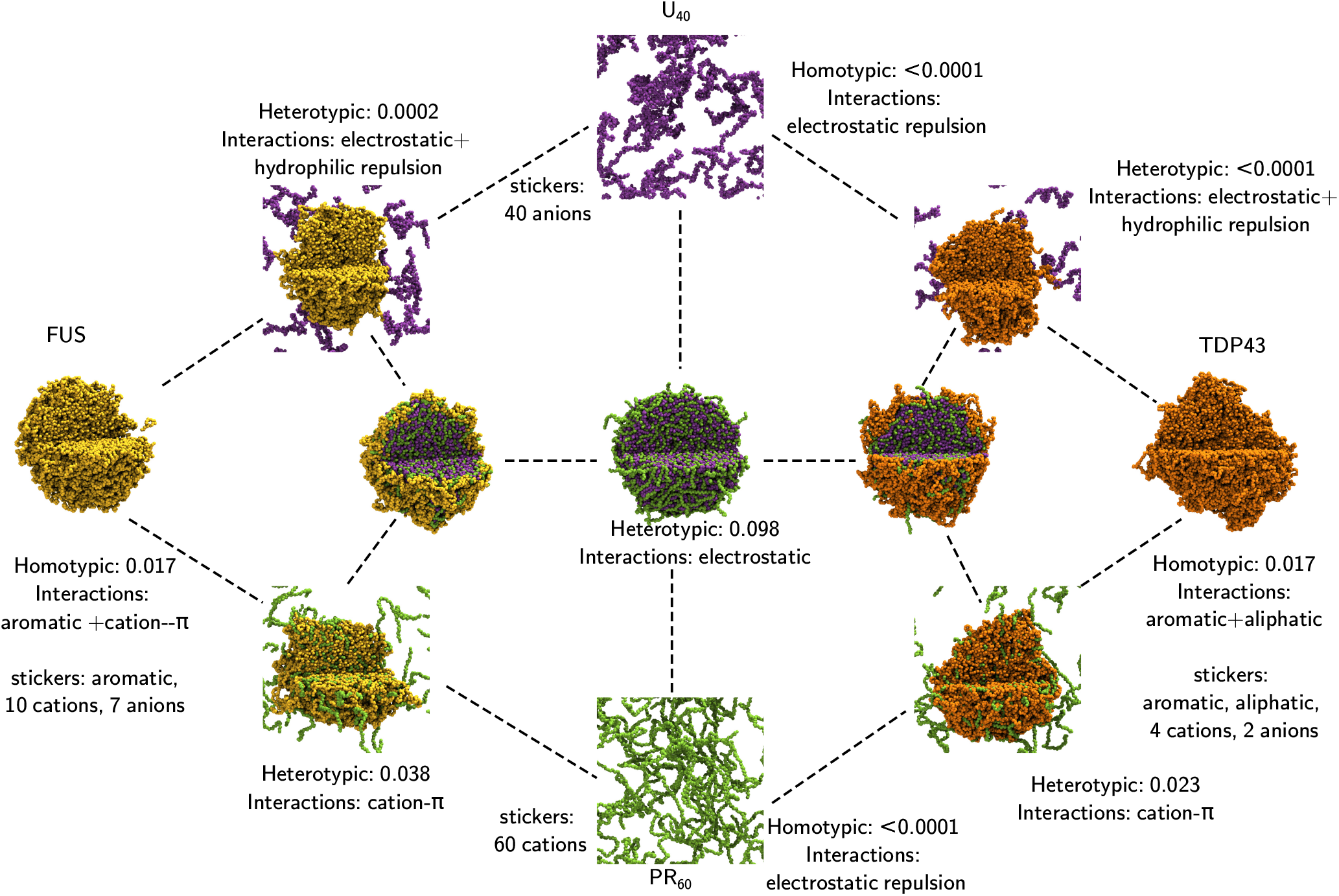
FUS-U_40_-PR_60_ and TDP43-U_40_-PR_60_ interaction triangles.

In simulations of ternary compositions of TDP43 with U_40_ and R-DPRs the same behaviour as with FUS was observed, as shown in Fig. 5 (additional images of TDP43 with other R-DPRs ca nbe found in Fig. S3 in the SI). Here we see a symmetry in the behaviour, with FUS PLD+RGG1 or TDP43 CTD forming a shell-like coating on the surface of a U_40_-PR_60_ droplet. Such behaviour indicates a potential universal mechanism for R-DPR induced toxicity that involves the sequestration of RNA with the RBPs forming coatings on the surface, providing a suitable interface for aggregation.

### Aliphatic interactions drive length dependent LLPS of ΔR-DPRs

The ΔR-DPRs have no arginine to drive intermolecular interactions with RNA and RBPs, leaving the aliphatic residue (proline or alanine) in the DPR to be the only residue to enable LLPS. Fig. S4 shows that self condensation of the ΔR-DPRs is length dependent with GA_15_ being too short to enable phase separation, whereas GA_30_, GA_60_, GP_60_ and PA_60_ do form a droplet. Upon addition of U_40_ (second row of Fig. S4) no change in ΔR-DPR LLPS behaviour is observed. This is due to the repulsive hydrophilic interactions of the RNA phosphate group with the aliphatic residues in the ΔR-DPRs.

Interestingly, the addition of RBPs does lead to changes in condensate behaviour. FUS and TDP43 both undergo LLPS in the presence of ΔR-DPRs. For GA_15_, the shortest ΔR-DPR studied, FUS and TDP43 condensates show no significant interactions with the ΔR-DPR, resulting in GA_15_ remaining in the dilute phase. For the longer ΔR-DPRs, condensate formation still occurs for both and significant interactions between the two condensates are observed. FUS and TDP43 form bimodal droplets with GA_30_, GA_60_ and GP_60_, and a coated droplet with the more hydrophobic PA_60_ as shown in Fig. S4. This behaviour is significantly stronger for TDP43 which has a higher aliphatic residue contact allowing for stronger heterotypic interactions that results in full covering of the PA_60_ droplet, whereas only partial coverage is observed for FUS. Interpenetration of the droplets is not observed, since the heterotypic interactions are weaker than the homotpyic interactions (see Fig. 6).

**Figure 6:**
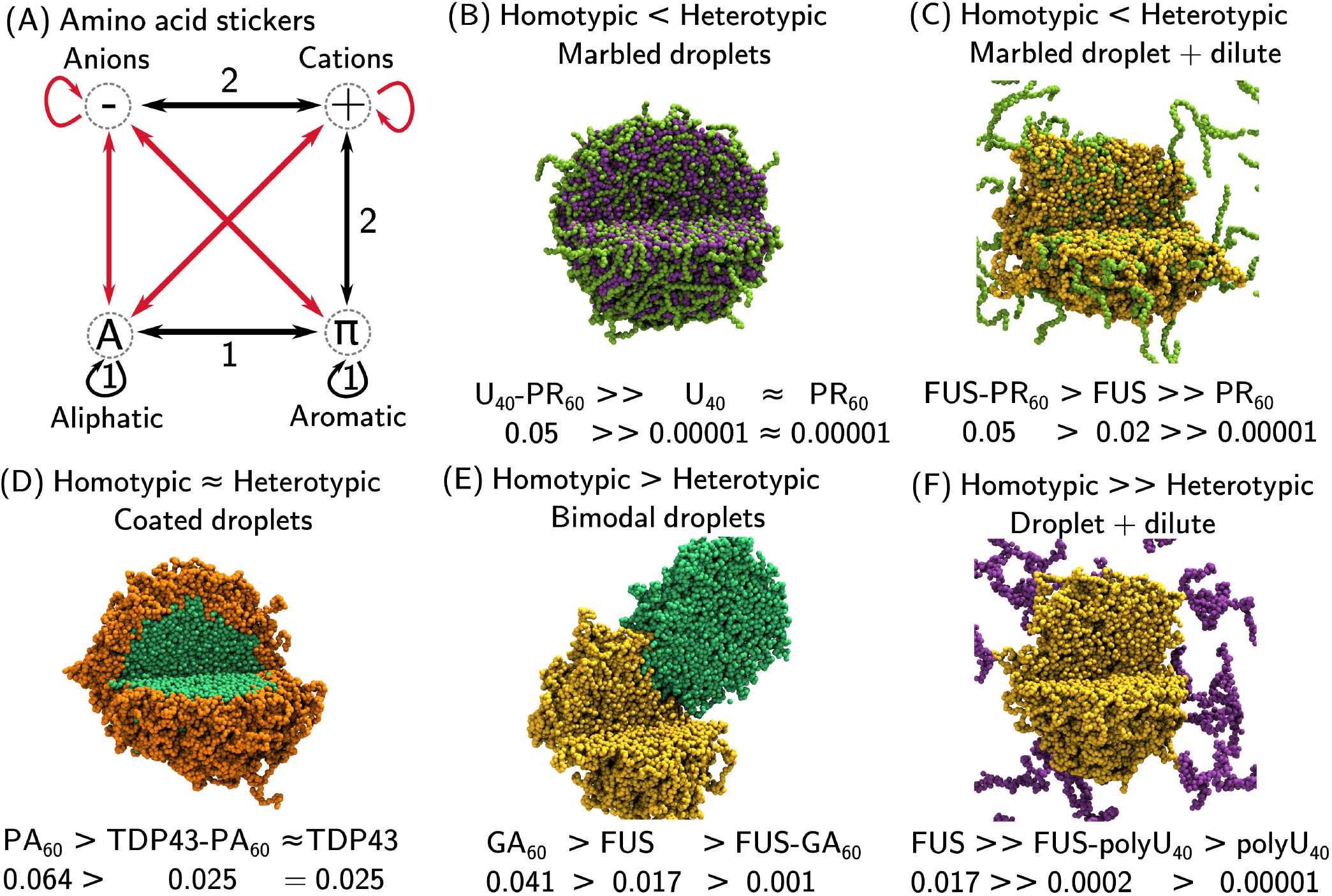
Summary of condensate interactions. (A) Amino acid sticker types and interactions. Favourable interactions (black) promote condensate formation, unfavourable interactions (red) oppose condensate formation. Sticker interactions fall into two orthogonal groupings: 1) Hydrophobic interactions (aliphatic, aromatic and alipatic-aromatic contacts), and 2) Cationic interactions (Cation-*π* and electrostatic). (B)-(F) The five cases of relative homotypic and heterotypic interaction strengths in (50%-50%) mixtures that result in different condensate morphologies, illustrated with examples from simulations undertaken in this work.

## DISCUSSION

The LLPS of FUS and TDP43 are both driven by hydrophobic interactions. In FUS this is from aromatic (tyrosine) interactions in the PLD, whereas aliphatic interactions provide the largest contribution in TDP43 LLPS. Interestingly, these interactions are orthogonal to the electrostatic interactions that drive association with RNA (Fig. 6A). This is shown by the arginine-phosphate interactions of the FUS RGG1 domain (an RNA binding domain) with U_40_ (Fig. 2D). The repulsive interactions with the PLD domain provide poor solubility of U_40_ in FUS PLD+RGG1 droplets such that RNA is restricted to interact with the condensate surface. The analogous behaviour of TDP43 indicates that RBP LLPS driven by IDRs with low charge produce condensates with poor RNA solubility, restricting RNA to interact with any RNA binding motif that is exposed on the condensate surface. This is supported by evidence of the effect of charge patterning on the LAF1 IDR in the work of Regy *et al*. (62), where charge partitioning results in exclusion of A_15_ from the core of the LAF1 condensate.

R-DPRs are able to induce the formation of complex coacervates with RNA through strong electrostatic interactions. R-DPRs are also able to form cation-*π* interactions with both FUS and TDP43, leading to inclusion into the RBP condensate (Figs. 5 and 6C). The presence of R-DPRs therefore changes the observed LLPS behaviour of FUS/TDP43 with U_40_ through these strong heterotypic interactions. This strong interaction of R-DPRs is postulated to be the driving force for their toxicity. Fig. 7A and C show the formation of marbled U_40_ and PR_60_ coated by a layer of FUS or TDP43, respectively, whereas in the absence of PR_60_ the U_40_ remains in the dilute phase with only transient interactions with the FUS or TDP43 condensate. Recent evidence suggests that the interfaces of condensates are incredibly important sites for nucleation and growth of aggregates or fibrils of misfolded proteins (63–65). The increased surface area of RBP condensates from the formation of core-shell structures could therefore provide such a mechanism to increase the rate of aggregation transitions in cells where R-DPRs are expressed. The observed length dependence on the ternary phase diagram of FUS-U_40_-R-DPR provides support for this theory. The increased multivalency of longer R-DPRs results in the increased strength of RNA interaction and growth of central (white) region in the phase diagrams (Figs. 3 and S2). The white region is characterised by the formation of FUS coated U_40_-R-DPR droplets leading to FUS exposed on the outer condensate surface, that could provide new nucleation sites for misfolding. The coarse-grained simulations showed no discernable difference in the behaviour of PR or GR R-DPRs.

**Figure 7:**
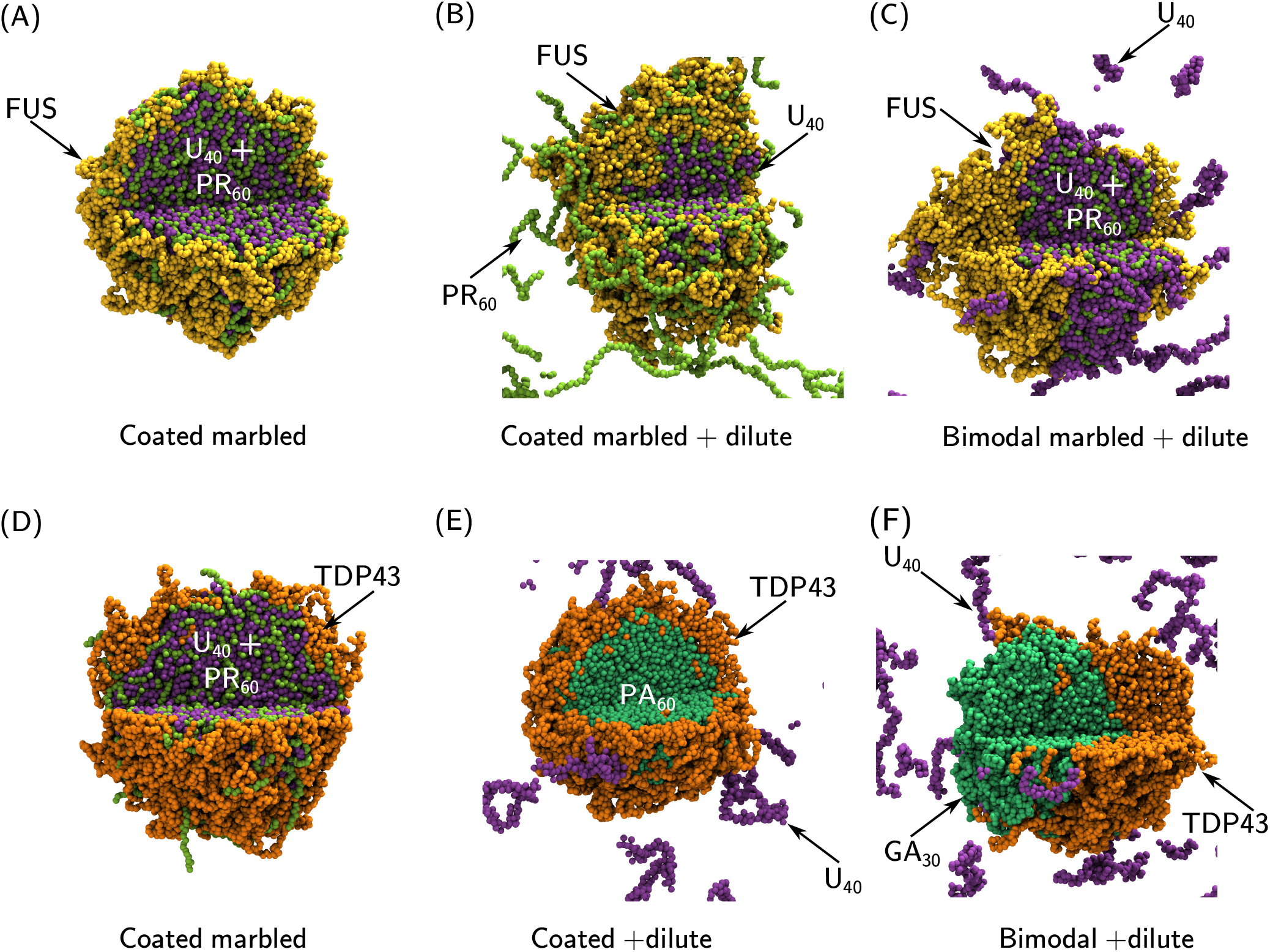
Simulation images for ternary mixtures showing the range of observed morphologies. (A) FUS-U_40_-PR_60_ (33.3, 33.3, 33.3), (B) FUS-U_40_-PR_60_ (37.5, 12.5, 50), (C) FUS-U_40_-PR_60_ (37.5, 50, 12.5), (D) TDP43-U_40_-PR_60_ (33.3, 33.3, 33.3), (E) TDP43-U_40_-PA_60_ (33.3, 33.3, 33.3), (F) TDP43-U_40_-GA_30_ (33.3, 33.3, 33.3), All simulations were run with a total amino acid concentration of 80000 *μM*. The end frame of 3 *μ*s of simulation is displayed as a representative state. A segment is not displayed to reveal the internal droplet structure. FUS molecules are coloured in yellow, TDP43 molecules are coloured in orange, U_40_ molecules are coloured in purple, PR_60_ molecules are coloured in light green, PA_60_ molecules are coloured in green, and GA_30_ molecules are coloured in green.

In contrast to R-DPRs, ΔR-DPRs do not interact with RNA, and short ΔR-DPRs do not interact with the IDRs of the RBPs FUS and TDP43 thus providing no effect on RBP LLPS in correspondence with experimental observations (24, 30, 32, 52, 53). Longer ΔR-DPRs do possess the ability to form condensates through hydrophobic aliphatic interactions which allow for interaction with RBP condensates through hydrophobic contacts. Such heterotypic interactions of ΔR-DPRs with FUS and TDP43 are significantly weaker than those of the R-DPRs with FUS and TDP43, resulting in coated droplet or bimodal droplet morphologies (shown in Fig. 6D and E). The TDP43-ΔR-DPR interactions do not influence the TDP43-RNA interactions in ternary systems (Fig. 7E and F), so therefore we do not expect the same mechanism for toxicity. Previous *in vivo* studies of ΔR-DPRs found that GA is able to aggregate and, interestingly, is required to enable aggregation of PA and GP ΔR-DPRs (52, 53). This indicates that the proline in GP and PA provides a barrier to the misfolding and aggregation of ΔR-DPRs. The interactions of GA ΔR-DPR condensates with TDP43 condensates forming bimodal droplets provides a potential pathway for the increased rates of TDP43 cleavage (52, 53). This is hypothesised to be caused by the interaction of the GA condensates with the cleavage enzymes, with co-localisation on the GA droplet surface enabling increased cleavage rates on TDP43 in interacting droplets.

## MATERIALS AND METHODS

### Protein and RNA sequences

We use FUS PLD+RGG1 domains (residues 1-267) in our simulations. The PLD (prion like domain, residues 1-167) is known to drive LLPS (66), with the RGG1 (arginine-glycine-glycine repeat domain, residues 168-267) enabling RNA interactions. The C terminal IDR (residues 273-414) was used to represent TDP43. U_40_ RNA was used as the model RNA strand in the simulations. The DPR proteins with repeating units PR, GR or GA containing either 60, 30, or 15 repeats, and DPR proteins with repeating units PA and GP containing 60 repeats were used (11 different DPR sequences in total). These repeat lengths were chosen to represent expansions observed in healthy individuals (15 repeats), transition individuals (30 repeats), and disease individuals (60 repeats).

### The 1 bead per amino acid (1BPA) molecular dynamics model

The 1BPA model was previously developed for the study of intrinsically disordered proteins in the nuclear pore complex (NPC) (67, 68). The 1BPA model used in this work uses the updated parameters described in (69).

### The 3 bead per nucleic acid (3BPN) molecular dynamics model

The 3BPN model for disordered RNA was designed to provide a 1BPA compatible model to be able to simulate mixtures of IDPs and ssRNA. It is based on the three interaction site model of RNA (70). 1BPA-compatible hydrophobicities were generated by first computing the Kapcha-Rossky hydrophobicity (71) of the 3BPN beads and subsequently using a linear mapping to the 1BPA hydrophobicities of the amino acids (see SI).

### Droplet simulation protocol

To be able to explore the behaviour of the transcription condensates we used droplet formation simulations to be able to look at the internal structure. Self-assembly and clustering of individual monomers into phase separated condensates can be a slow process to observe. To speed up this process we start by forming a condensed phase droplet at the start of the simulation, which is then inserted into an empty dilute phase. If LLPS is favoured, this droplet structure should remain stable throughout the subsequent simulation; if unstable this droplet would breakup into a dilute phase of monomers.

The initial cubic simulation box is populated with molecules (using a random initial conformation) with their center of mass placed upon a regular grid, with a small buffer region to avoid overlap between molecules. All simulations are carried out at a temperature of 300 K and use a timestep of 20 fs. For equilibration of the droplet, energy minimisation on the initial configuration is used (energy tolerance of 1 kJ mol^−1^ nm^−1^), before 50 ns NVT Langevin dynamics simulations (Nosé-hoover thermostat with *τ*_*t*_ = 100 ps), followed by 500 ns NPT Langevin dynamics (Nosé-hoover thermostat with *τ*_*t*_ = 100 ps and a Berendsen barostat with *τ*_*p*_ = 10 ps, 1 bar reference pressure and a compressibility of 4.5 ×10^−5^ bar^−1^). The end state of the NPT equilibration step is inserted into a new periodic box with a volume chosen to give a total particle density of 80,000 *μ*M, after recentering on the center of mass and after the molecules have been unwrapped across the previous periodic boundary conditions. A second energy minimisation step is applied in the new simulation box to relax the molecules after the box expansion (energy tolerance of 1 kJ mol^−1^ nm^−1^). A final 3 *μ*s NVT production run (Nosé-hoover thermostat with *τ*_*t*_ = 100 ps) is used for data collection. The trajectory is sampled every 5 ns to determine whether convergence was reached.

## AUTHOR CONTRIBUTIONS

MDD and PRO designed the research. MDD and JP carried out all simulations. MDD analyzed the data. MDD wrote the first draft. JP reviewed the article. PRO reviewed and edited the article and supervised the research.

## ACKNOWLEDGMENTS

We thank the oLife COFUND project for funding MDD. The COFUND project oLife has received funding from the European Union’s Horizon 2020 research and innovation programme under grant agreement No 847675. We thank the Center for Information Technology of the University of Groningen for their support and for providing access to the Peregrine high performance computing cluster. This work made use of the Dutch national e-infrastructure with the support of the SURF Cooperative using grant no. EINF-3456 and EINF-5917.

## SUPPLEMENTARY MATERIAL

Supplementary information contains the FUS and TDP43 protein sequences, 3BPN model details, additional simulation images and ternary phase diagrams for FUS and U_40_ with additional R-DPR sequences.

## Notes

### Competing Interest Statement

The authors have declared no competing interest.

