## supplementary information for "The effect of dipeptide repeat proteins on FUS/TDP43-RNA condensation in C9orf72 ALS/FTD"

M.D. Driver

May 21, 2024

#### Contents


**1 1BPA model** **2**

**2 3BPN model** **2**

**3 Simulation analysis using a molecular connectivity graph** **3**

3.1 Readouts from a molecular connectivity graph . . . . . . . . . . . . . . . . . . . . . . . . . . . . 3

3.2 Contact map definition . . . . . . . . . . . . . . . . . . . . . . . . . . . . . . . . . . . . . . . . . 4

**4 Ternary phase diagram comparison** **5**

**5 TDP43-R-DPR and ΔR-DPR binary mixture comparisons** **6**

**6 Droplet simulation data** **7**

### 1 1BPA model

The 1BPA model was previously developed for the study of the nuclear pore complex (NPC) [1, 2]. It has been extensively used to model the behaviour of the intrinsically disordered nucleoporin regions which fill the center of the NPC and provide a selective barrier to the transport of cargo between the cytoplasm and nucleoplasm [3–12]. In this work we use the updated 1BPA v2.1 developed in [13].

### 2 3BPN model

The 3 bead per nucleotide (3BPN) model was designed to provide a 1BPA compatible model to be able to simulate mixtures of IDPs and ssRNA. It is based on the three interaction site model of RNA [14].

The 3BPN bonded potential, $\phi_b$, is composed of bonds $\phi_{bond}$ and bending angles $\phi_{bend}$, which both take the form of simple harmonic potentials:

$$\phi_b = \phi_{bond} + \phi_{bend}, \tag{1}$$

with $\phi_{\mathrm{bond}}$ and $\phi_{\mathrm{bend}}$ given by,

$$\phi_{\mathrm{bond}} = k_b(r - r_0)^2, \tag{2}$$

$$\phi_{\mathrm{bend}} = k_\theta(\theta - \theta_0)^2, \tag{3}$$

with $r_0$ and $k_b$ given in Table S1 and $\theta_0$ and $k_\theta$ given in Table S2.

| Bonds | $r_0$ /nm | $k_b$ / kJ mol$^{-1}$nm$^{-2}$ |
|---|---|---|
| $NB_i$-$NR_i$ | 0.586 | 10000 |
| $NP_i$-$NR_i$ | 0.390 | 10000 |
| $NP_i$-$NR_{i+1}$ | 0.390 | 10000 |

Table S1: Bonded parameters where $NB_i$ is the nucleobase bead of nucleotide $i$ (representing either G, C, A or U), $NR_i$ is the ribose bead of nucleotide $i$ and $NP_i$ is the phosphate bead of nucleotide $i$. Values for $r_0$ and $k_b$ are taken from [14].

| Angles | $\theta_0$ / rad | $k_\theta$ / kJ mol$^{-1}$rad$^{-2}$ |
|---|---|---|
| $NB_i$-$NR_i$-$NP_i$ | 1.76 | 1000 |
| $NP_i$-$NR_{i+1}$-$NB_{i+1}$ | 1.59 | 1000 |
| $NR_i$-$NP_i$-$NR_{i+1}$ | 1.86 | 1000 |
| $NP_{i-1}$-$NR_i$-$NP_i$ | 1.97 | 1000 |

Table S2: Angle parameters where $NB_i$ is a nucleotide bead of nucleotide $i$ (representing either G, C, A or U), $NR_i$ is a ribose bead of nucleotide $i$, $NP_i$ is a phosphate bead of nucleotide $i$. Values of $\theta_0$ and $k_\theta$ are taken from [14].

There are three components that constitute the non-bonded potential, $\phi_{nb}$ (equation (4)), of the 3BPN model: hydrophobic interactions ($\phi_{hp}$, equation (5)), base stacking interactions ($\phi_{bs}$, equation (6)) and electrostatic interactions ($\phi_{el}$, equation (7)). The hydrophobic (equation (5)) and electrostatic terms (equation (7)) are equivalent to the 1BPA hydrophobic and electrostatic terms to ensure cross compatibility between the two models and enable the simulation of protein-RNA mixtures. Also the $\phi_{bs}$ has been adjusted to have a similar funtional form as the 1BPA hydrophobicity potential.

$$\phi_{nb} = \phi_{hp} + \phi_{bs} + \phi_{el}, \tag{4}$$

$$\phi_{hp} = \begin{cases} \epsilon_{rep}(\frac{\sigma}{r})^8 - \epsilon_{ij}[\frac{4}{3}(\frac{\sigma}{r})^6 - \frac{1}{3}], & r \leq \sigma \\ \epsilon_{rep} - \epsilon_{ij}(\frac{\sigma}{r})^8 & \sigma \leq r, \end{cases} \tag{5}$$

where $\epsilon_{ij} = \epsilon_{hp}\sqrt{(\epsilon_i \epsilon_j)^\alpha}$, $\sigma = 0.6$ nm, $\epsilon_{rep} = 5$ kJ mol$^{-1}$, $\alpha = 0.15$, $\epsilon_{hp} = 6.5$ kJ mol$^{-1}$ and $\epsilon_i$, $\epsilon_j$ are bead specific hydrophobicities. These were determined by using the Kapcha-Rossky hydrophobicity scale [15] with the 1BPA amino acid hydrophobicities [1, 2], to create a mapping for the nucleotide beads between the Kapcha-Rossky hydrophobicity and the corresponding 1BPA hydrophobicity, see Fig. S1 and the $\epsilon_i$ are given in Table S3.

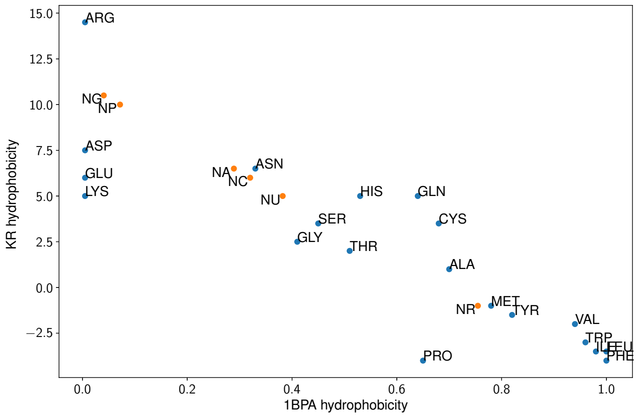


Figure S1: Hydrophobicity scale comparison for the Kapcha-Rossky hydrophobicity scale [15] with the 1BPA amino acid hydrophobicities [1, 2]. Amino acid hydrophobicities are shown by blue dots with the three letter amino acid name. A linear mapping was used to determine the corresponding nucleotide bead 1BPA hydrophobicities (orange dots, labelled by bead name) after computing the Kapcha-Rossky hydrophobicity.

| Nucleotide bead | $\epsilon_i$ |
|---|---|
| NP | 0.071 |
| NR | 0.755 |
| NA | 0.289 |
| NC | 0.320 |
| NG | 0.040 |
| NU | 0.382 |

Table S3: $\epsilon_i$ values for 3BPN nucleic acid beads.

$$\phi_{bs} = \begin{cases} \epsilon_{bs1}(\frac{\sigma}{r})^8 - \epsilon_{bs2}[\frac{4}{3}(\frac{\sigma}{r})^6 - \frac{1}{3}], & r \leq \sigma \\ \epsilon_{bs1} - \epsilon_{bs2}(\frac{\sigma}{r})^8 & \sigma \leq r, \end{cases} \tag{6}$$

where $\sigma = 0.6$ nm, $\epsilon_{bs1} = 2.71$ kJ mol$^{-1}$ and $\epsilon_{bs2} = 1.25$ kJ mol$^{-1}$.

$$\phi_{el} = \frac{q_i q_j}{4\pi\epsilon_0\epsilon_r(r)r} exp(-\kappa r), \tag{7}$$

where $\epsilon_r(r) = S_s[1 - \frac{r^2}{z^2}\frac{e^{r/z}}{(e^{r/z}-1)^2}$, $S_s = 80$ and $z = 0.25$ nm. $\kappa$ is the Debye screening coefficient which is a function of temperature and ion concentration.

### 3 Simulation analysis using a molecular connectivity graph

To assess condensate stability in the droplet simulations, a molecular connectivity graph, consisting of nodes and edges, is used to describe the clustering and assess the time evolution of the molecular network during the simulation. Each simulation frame can be described by a connectivity graph. In the graph representation each node corresponds to a unique molecule, with an edge between the nodes denoting the number of non-zero interactions (if zero interactions are present no edge is added). Within an edge information can be stored about the number and type of contacts between molecules, based on residue categorisation. An interaction was determined to be present using a cutoff of 0.7 nm between 1BPA residues. Computation and processing of the contact matrix for a simulation frame allows the creation of the molecule graph. This process is shown in figure S2.

#### 3.1 Readouts from a molecular connectivity graph

A molecular connectivity graph computed for a simulation can be used to get several different readouts about a simulation system. Information on the size of clusters can be found by identifying the discrete sub-graphs of nodes with only internal connections (no contacts outside of the cluster). Through assessment

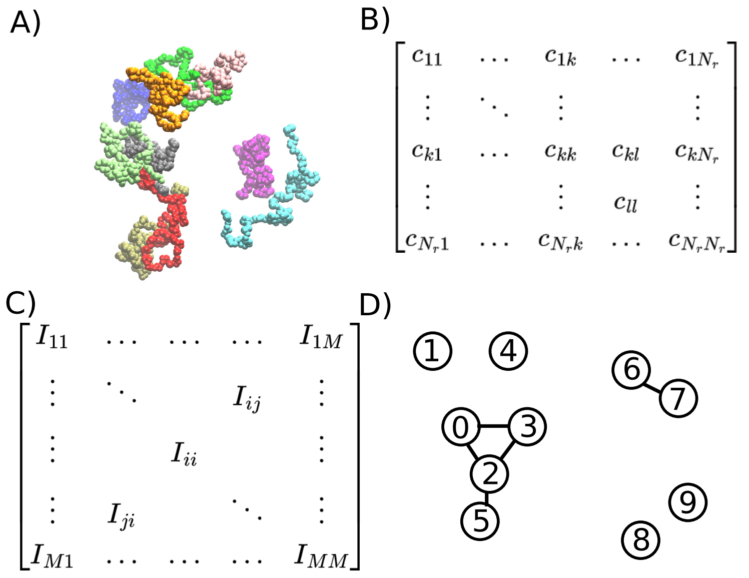


Figure S2: Processing of a contact matrix for one trajectory frame. (A) Visualisation of molecules in the trajectory frame. The particles of the 10 molecules are coloured based on the molecule type. (B) The contact matrix for a frame is computed between all residues, yielding an $N_r \times N_r$ matrix, with a value of 1 if a contact is present between particles $k$ and $l$, and 0 otherwise. (C) The contact matrix in (B) can be decomposed into distinct sub-matrices by grouping together residue interactions belonging to unique molecule pairs. This produces a new matrix representation, where element $I_{ij}$ is the contact matrix between molecules $i$ and $j$. (D) A molecular graph derived from the contact matrix in (C). When constructing the molecular graph we consider only the elements where $i \geq j$ to avoid double counting of interactions. We iterate over the $I_{ij}$ matrices ($i \neq j$), to add edges to the graph, storing information about the number of intermolecular contacts between molecules $i$ and $j$. Similarly, a summation of intramolecular contacts $I_{ii}$ for like molecules (case when $i = j$) is also stored. This process is repeated for all frames in a simulation with a sampling rate of once every 50 ns.

of the time evolution of the number of clusters, grouped based on number of molecules, and the number of molecules in clusters within specific size categories it is possible to determine whether the system has reached equilibrium. After convergence is established it is possible to examine the distribution of components within the different clusters. The percentage of species $i$ in a cluster is computed using $N_{i\in C}/(N_{frames}N_i)$ where $N_{i\in C}$ is the number of molecules of $i$ in a cluster of size $C$.

In addition to collating node information from a cluster, the number and type of interactions stored in the graph edges can also be used to assess the simulations. Interaction data can be aggregated based on the type of species involved. Normalisation is again required to be able to compare the relative number of interactions between molecules, using $N_iL_iN_jL_j$, where $L_i$ and $L_j$ are the number of simulation residues in molecules $i$ and $j$, respectively.

##### 3.2 Contact map definition

From the system graph creation process, contact matrices for all molecule combinations, over all frames are computed. This information is also useful for understanding the specific residues which drive condensation. As such, this information is aggregated during graph computation. Contact maps can be divided into two categories: intramolecular contact maps (interactions between particles in the same molecule copy), corresponding to the diagonal sub-matrices ($I_{ii}$) in figure S2, and intermolecular contact maps (interactions between particles of two different molecules), corresponding to the off-diagonal sub-matrices ($I_{ij}$) in figure S2. On the macroscale individual molecules of the same type are indistinguishable, thus only unique combinations of molecules need to be stored in different matrices. A system has $M$ intramolecular contact maps and $M(M+1)/2$ intermolecular contact maps where $M$ is the number of distinguishable species. Aggregation over all frames and molecule copies of the contact maps is therefore possible for memory efficient storage during computation. To display the information stored in the contact map as a contact probability, ordered by particle index in a molecule, the contact information is normalised by the number of frames and $((N_iN_j)^2)$ where $N_i$ and $N_j$ are the number of copies of molecule types $i$ and $j$ in the simulation. This is to allow comparison between species in the simulation where the number of copies may be very large. To display the information based on residue type a matrix reduction is used on the contact map plotted by residue index to sum interactions of particles with the same type. Further normalisation can be undertaken of this matrix to normalise the interactions by $(N_{i,r1}N_{j,r2})^2$, with $N_{j,r1}$ the number of residue $r1$ in molecule $i$ and $N_{j,r2}$ the number of residue $r2$ in molecule $j$.

### 4 Ternary phase diagram comparison

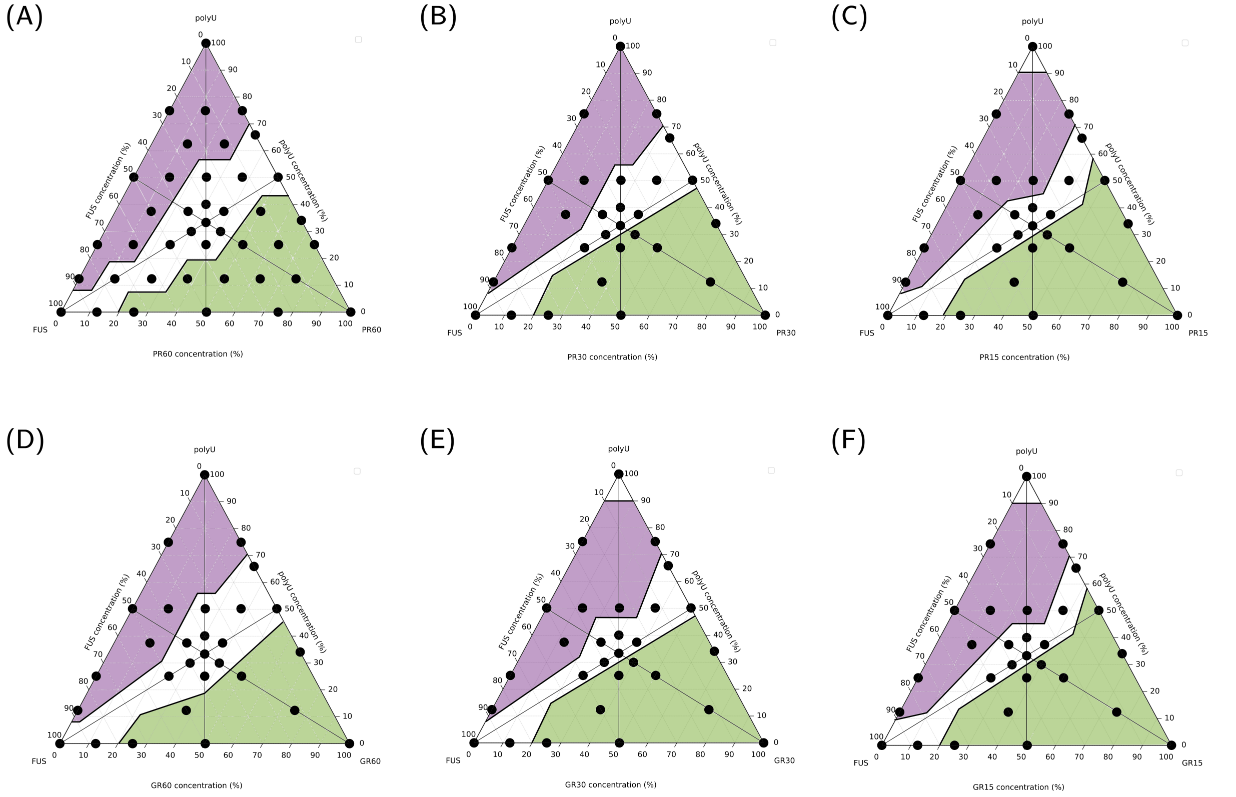


Figure S3: Ternary phase diagrams for (A) $PR_{60}$, (B) $PR_{30}$, (C) $PR_{15}$, (D) $GR_{60}$, (E) $GR_{30}$ and (F) $GR_{15}$. Triangles showing summary of phases found to highlight any changes- these are currently in SI for discussion. There is a small change to the two boundaries narrowing on the centre region with R-DPR length.

### 5 TDP43-R-DPR and ΔR-DPR binary mixture comparisons

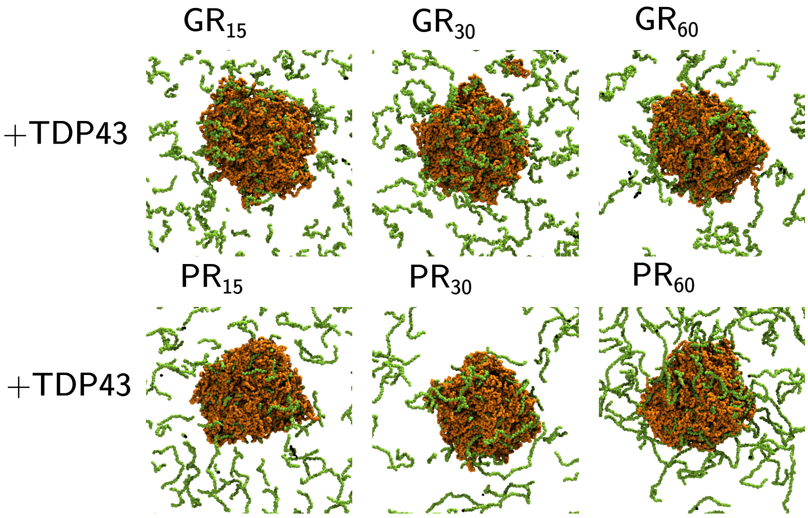


Figure S4: Figure showing the behaviour of TDP43 with different R-DPR sequences. All simulations are done at equal fraction DPR to TDP43.

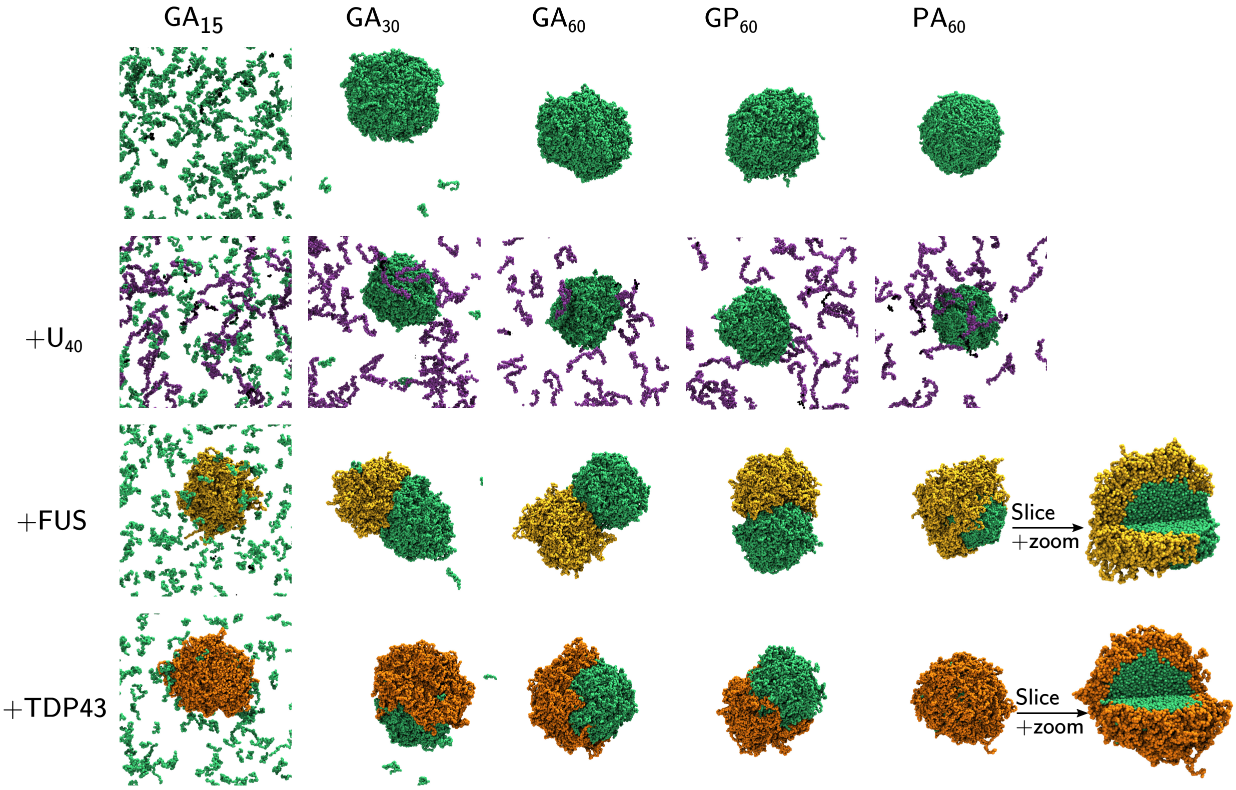


Figure S5: PS behaviour of ΔR-DPR sequences in isolation and with various partners. Top row isolated ΔR-DPR, second row ΔR-DPR with $U_{40}$, third row ΔR-DPR with FUS, fourth row ΔR-DPR with TDP-43. All simulations are done at an equal fraction of DPR to $U_{40}$/FUS/TDP43.

### 6 Droplet simulation data

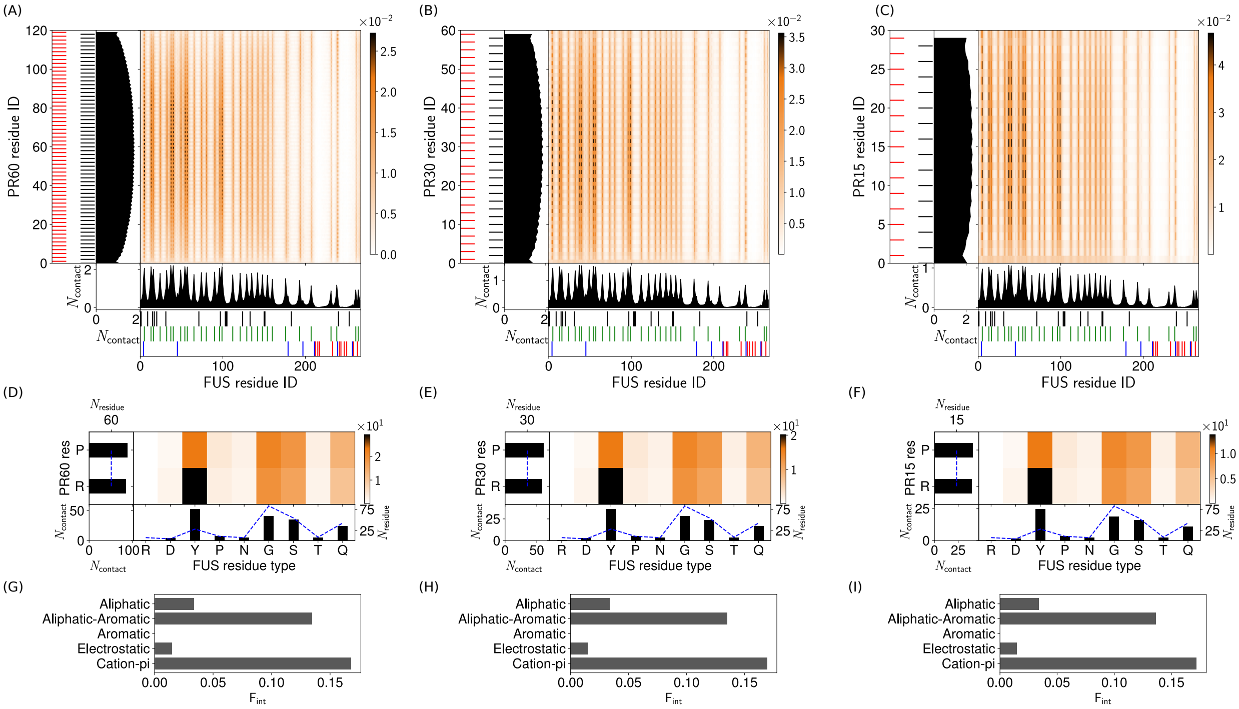

Figure S6: **Intermolecular contact maps for heterotypic interactions in two-component droplets.** (A)-(C) Intermolecular contact map by residue index for (A) FUS with PR60 (50%, 50%), (B) FUS with PR30 (50%, 50%), and (C) FUS with PR15 (50%, 50%) at 150 mM and 300 K. For the definitions of the different contact types see section 3. The 1D contact profiles denote a summation of the 2D map of the corresponding molecules. The black dashed lines highlight the key residues: the aromatic residues in FUS. (D)-(F) Intermolecular contact map by residue type for (D) FUS with PR60 (50%, 50%), (E) FUS with PR30 (50%, 50%), and (F) FUS with PR15 (50%, 50%) at 150 mM and 300 K. (G)-(I) Intermolecular interaction summary for (G) FUS-PR60 interactions in (50%, 50%), (H) FUS-PR30 interactions in (50%, 50%), and (I) FUS-PR15 interactions in (50%, 50%) at 150 mM and 300 K. The fraction of interactions, $\mathsf{F}_{\mathsf{int}}$, are aggregated by type and normalised by the total number of the intermolecular interactions in (A)-(C) respectively.

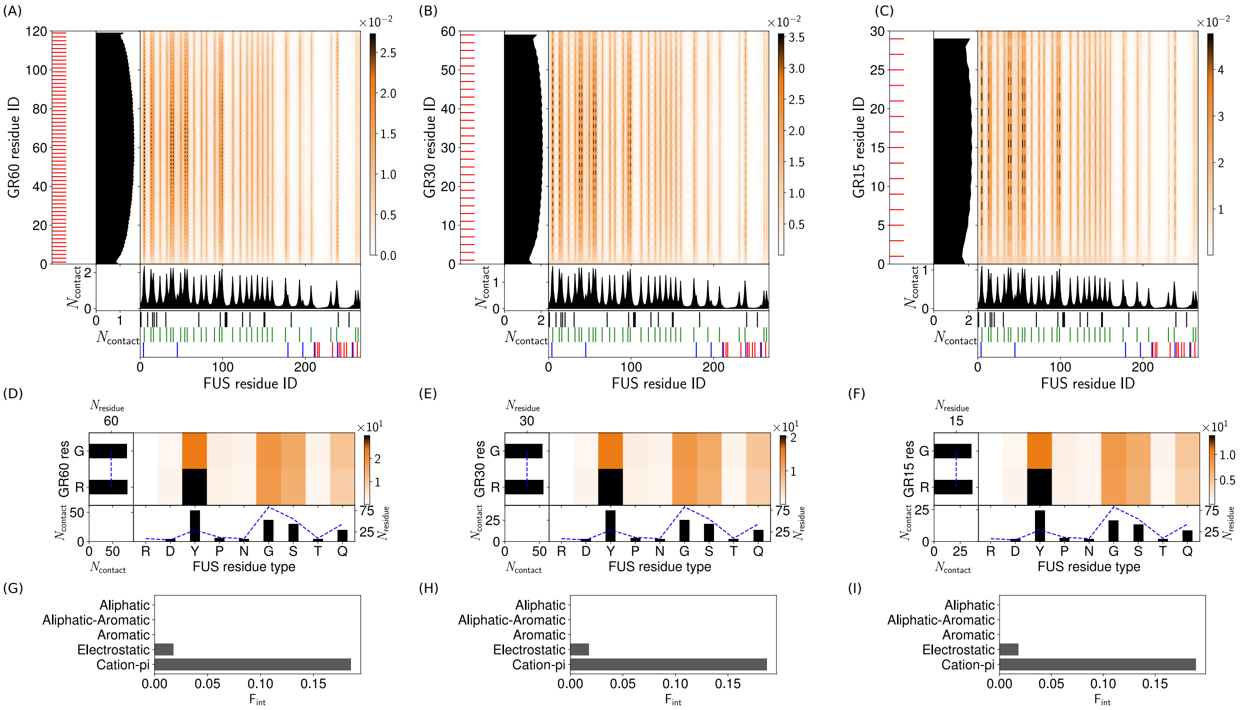


Figure S7: **Intermolecular contact maps for heterotypic interactions in two-component droplets.** (A)-(C) Intermolecular contact map by residue index for (A) FUS with GR60 (50%, 50%), (B) FUS with GR30 (50%, 50%), and (C) FUS with GR15 (50%, 50%) at 150 mM and 300 K. For the definitions of the different contact types see section 3. The 1D contact profiles denote a summation of the 2D map of the corresponding molecules. The black dashed lines highlight the key residues: the aromatic residues in FUS. (D)-(F) Intermolecular contact map by residue type for (D) FUS with GR60 (50%, 50%), (E) FUS with GR30 (50%, 50%), and (F) FUS with GR15 (50%, 50%) at 150 mM and 300 K. (G)-(I) Intermolecular interaction summary for (G) FUS-GR60 interactions in (50%, 50%), (H) FUS-GR30 interactions in (50%, 50%), and (I) FUS-GR15 interactions in (50%, 50%) at 150 mM and 300 K. The fraction of interactions, $\mathsf{F}_{\mathsf{int}}$, are aggregated by type and normalised by the total number of the intermolecular interactions in (A)-(C) respectively.

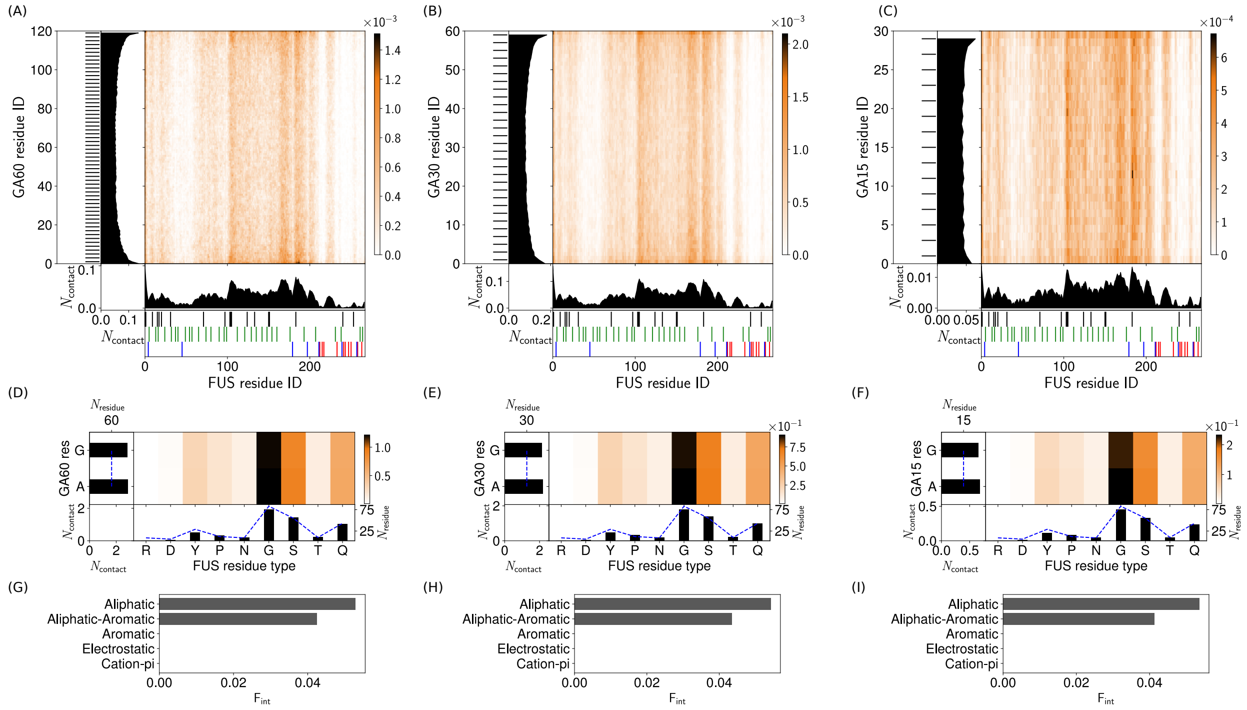


Figure S8: **Intermolecular contact maps for heterotypic interactions in two-component droplets.** (A)-(C) Intermolecular contact map by residue index for (A) FUS with GA60 (50%, 50%), (B) FUS with GA30 (50%, 50%), and (C) FUS with GA15 (50%, 50%) at 150 mM and 300 K. For the definitions of the different contact types see section 3. The 1D contact profiles denote a summation of the 2D map of the corresponding molecules. The black dashed lines highlight the key residues: the aromatic residues in FUS. (D)-(F) Intermolecular contact map by residue type for (D) FUS with GA60 (50%, 50%), (E) FUS with GA30 (50%, 50%), and (F) FUS with GA15 (50%, 50%) at 150 mM and 300 K. (G)-(I) Intermolecular interaction summary for (G) FUS-GA60 interactions in (50%, 50%), (H) FUS-GA30 interactions in (50%, 50%), and (I) FUS-GA15 interactions in (50%, 50%) at 150 mM and 300 K. The fraction of interactions, $F_{int}$, are aggregated by type and normalised by the total number of the intermolecular interactions in (A)-(C) respectively.

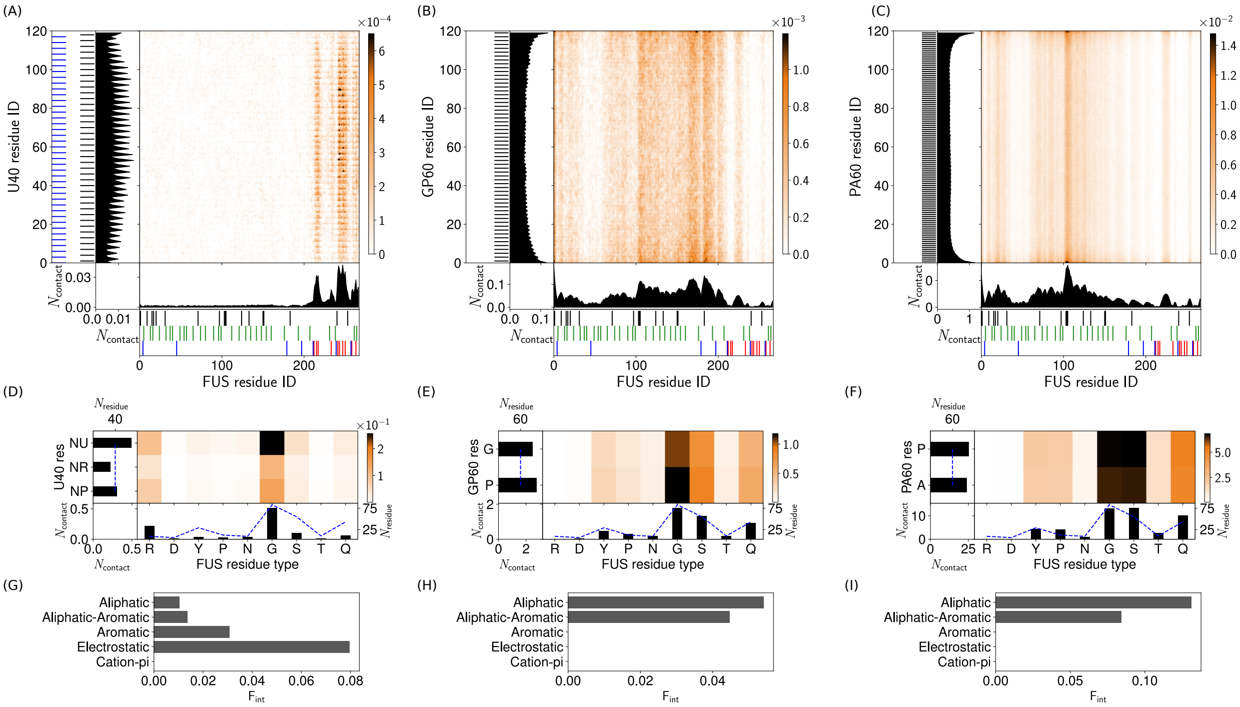


Figure S9: **Intermolecular contact maps for heterotypic interactions in two-component droplets.** (A)-(C) Intermolecular contact map by residue index for (A) FUS with U40 (50%, 50%), (B) FUS with GP60 (50%, 50%), and (C) FUS with PA60 (50%, 50%) at 150 mM and 300 K. For the definitions of the different contact types see section 3. The 1D contact profiles denote a summation of the 2D map of the corresponding molecules. The black dashed lines highlight the key residues: the aromatic residues in FUS. (D)-(F) Intermolecular contact map by residue type for (D) FUS with U40 (50%, 50%), (E) FUS with GP60 (50%, 50%), and (F) FUS with PA60 (50%, 50%) at 150 mM and 300 K. (G)-(I) Intermolecular interaction summary for (G) FUS-U40 interactions in (50%, 50%), (H) FUS-GP60 interactions in (50%, 50%), and (I) FUS-PA60 interactions in (50%, 50%) at 150 mM and 300 K. The fraction of interactions, $F_{int}$, are aggregated by type and normalised by the total number of the intermolecular interactions in (A)-(C) respectively.

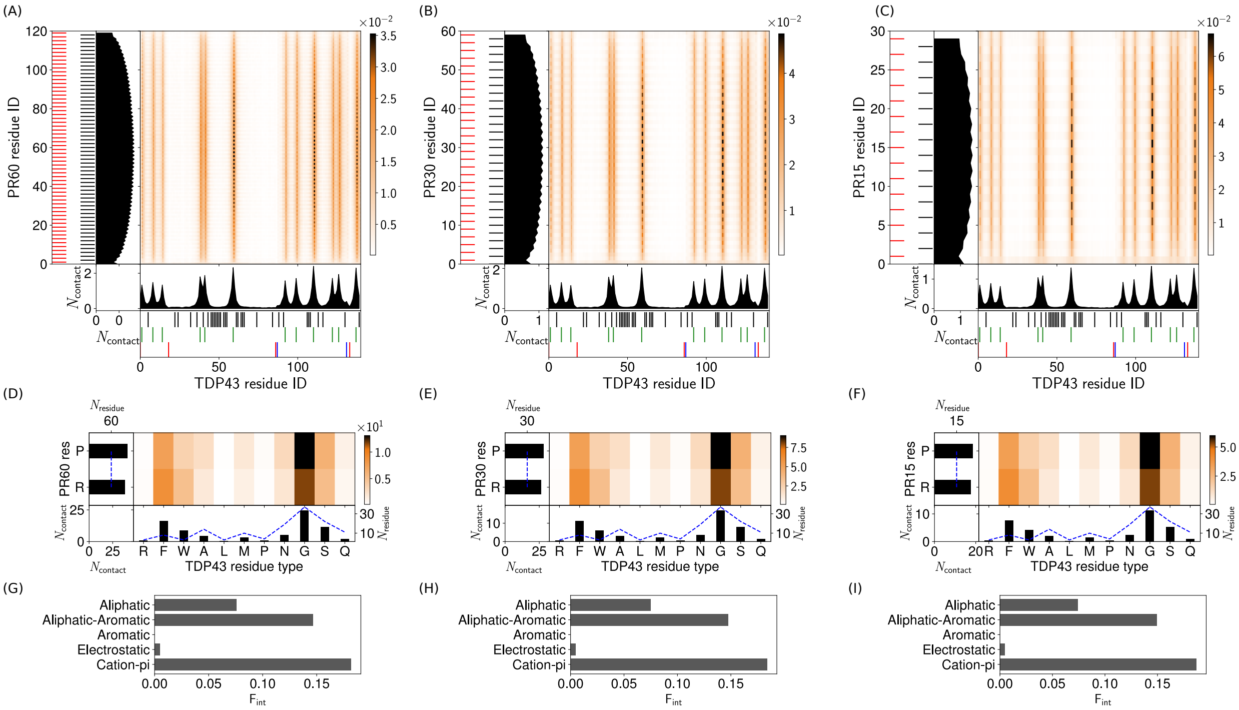


Figure S10: **Intermolecular contact maps for heterotypic interactions in two-component droplets.** (A)-(C) Intermolecular contact map by residue index for (A) TDP43 with PR60 (50%, 50%), (B) TDP43 with PR30 (50%, 50%), and (C) TDP43 with PR15 (50%, 50%) at 150 mM and 300 K. For the definitions of the different contact types see section 3. The 1D contact profiles denote a summation of the 2D map of the corresponding molecules. (D)-(F) Intermolecular contact map by residue type for (D) TDP43 with PR60 (50%, 50%), (E) TDP43 with PR30 (50%, 50%), and (F) TDP43 with PR15 (50%, 50%) at 150 mM and 300 K. (G)-(I) Intermolecular interaction summary for (G) TDP43-PR60 interactions in (50%, 50%), (H) TDP43-PR30 interactions in (50%, 50%), and (I) TDP43-PR15 interactions in (50%, 50%) at 150 mM and 300 K. The fraction of interactions, $\mathsf{F}_{\mathsf{int}}$, are aggregated by type and normalised by the total number of the intermolecular interactions in (A)-(C) respectively.

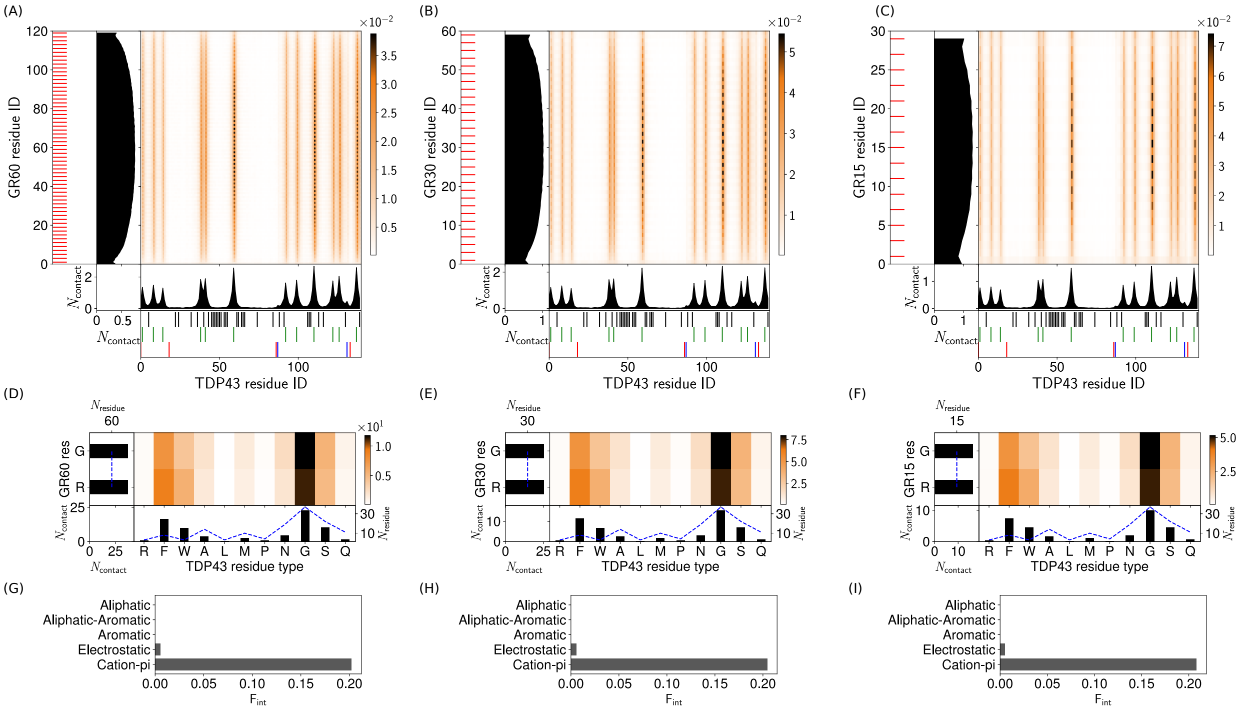

Figure S11: **Intermolecular contact maps for heterotypic interactions in two-component droplets.** (A)-(C) Intermolecular contact map by residue index for (A) TDP43 with GR60 (50%, 50%), (B) TDP43 with GR30 (50%, 50%), and (C) TDP43 with GR15 (50%, 50%) at 150 mM and 300 K. For the definitions of the different contact types see section 3. The 1D contact profiles denote a summation of the 2D map of the corresponding molecules. (D)-(F) Intermolecular contact map by residue type for (D) TDP43 with GR60 (50%, 50%), (E) TDP43 with GR30 (50%, 50%), and (F) TDP43 with GR15 (50%, 50%) at 150 mM and 300 K. (G)-(I) Intermolecular interaction summary for (G) TDP43-GR60 interactions in (50%, 50%), (H) TDP43-GR30 interactions in (50%, 50%), and (I) TDP43-GR15 interactions in (50%, 50%) at 150 mM and 300 K. The fraction of interactions, $\mathsf{F}_{\mathsf{int}}$, are aggregated by type and normalised by the total number of the intermolecular interactions in (A)-(C) respectively.

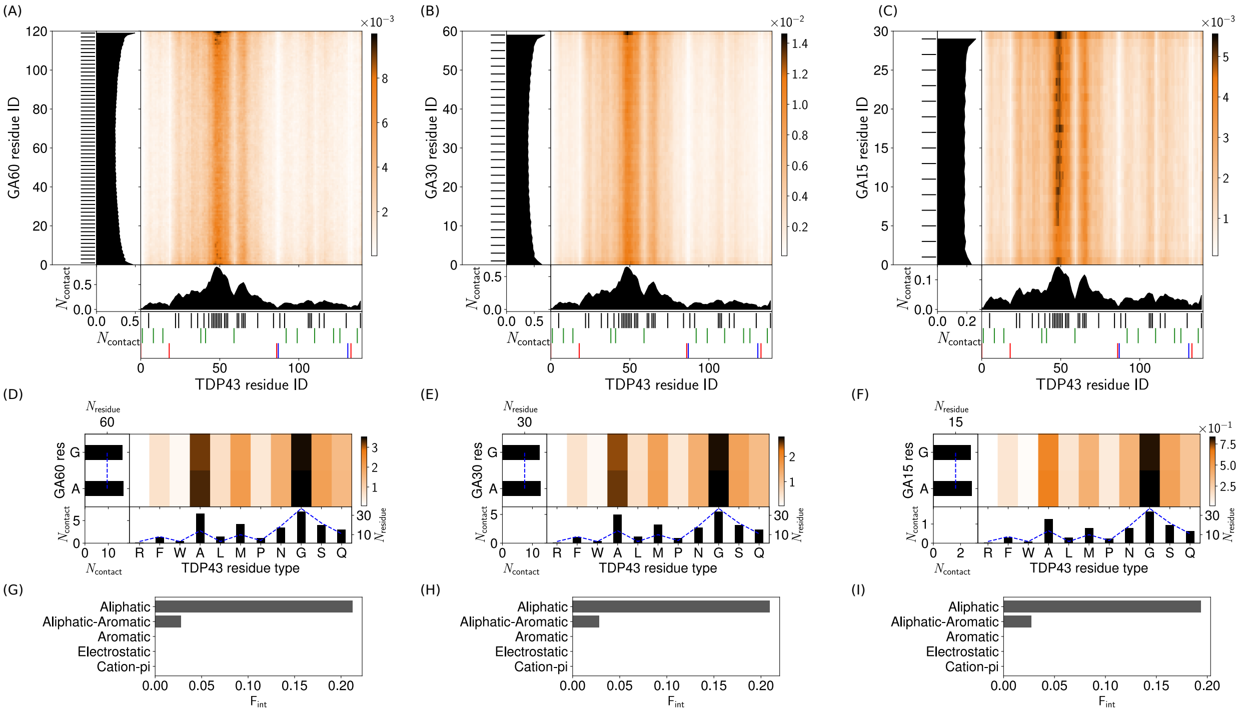


Figure S12: **Intermolecular contact maps for heterotypic interactions in two-component droplets.** (A)-(C) Intermolecular contact map by residue index for (A) TDP43 with GA60 (50%, 50%), (B) TDP43 with GA30 (50%, 50%), and (C) TDP43 with GAR15 (50%, 50%) at 150 mM and 300 K. For the definitions of the different contact types see section 3. The 1D contact profiles denote a summation of the 2D map of the corresponding molecules. (D)-(F) Intermolecular contact map by residue type for (D) TDP43 with GA60 (50%, 50%), (E) TDP43 with GA30 (50%, 50%), and (F) TDP43 with GA15 (50%, 50%) at 150 mM and 300 K. (G)-(I) Intermolecular interaction summary for (G) TDP43-GA60 interactions in (50%, 50%), (H) TDP43-GA30 interactions in (50%, 50%), and (I) TDP43-GA15 interactions in (50%, 50%) at 150 mM and 300 K. The fraction of interactions, $\mathsf{F}_{\mathsf{int}}$, are aggregated by type and normalised by the total number of the intermolecular interactions in (A)-(C) respectively.

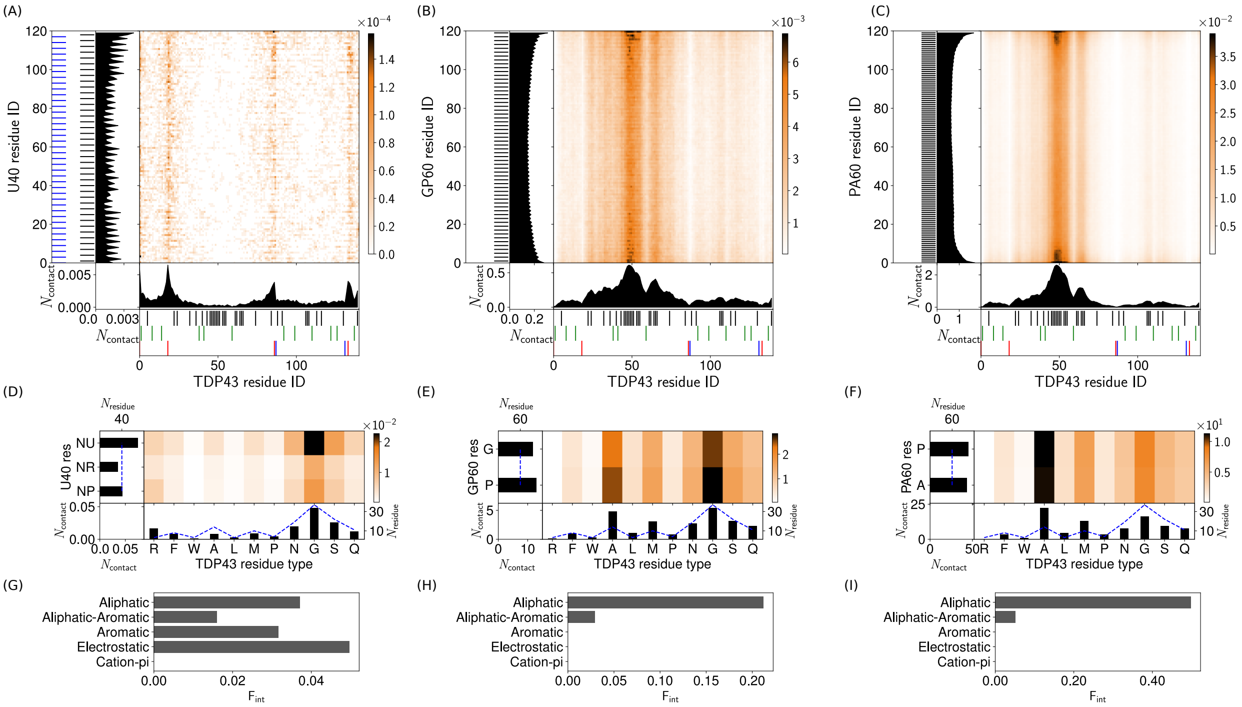


Figure S13: **Intermolecular contact maps for heterotypic interactions in two-component droplets.** (A)-(C) Intermolecular contact map by residue index for (A) TDP43 with U40 (50%, 50%), (B) TDP43 with GP60 (50%, 50%), and (C) TDP43 with PA60 (50%, 50%) at 150 mM and 300 K. For the definitions of the different contact types see section 3. The 1D contact profiles denote a summation of the 2D map of the corresponding molecules. (D)-(F) Intermolecular contact map by residue type for (D) TDP43 with U40 (50%, 50%), (E) TDP43 with GP60 (50%, 50%), and (F) TDP43 with PA60 (50%, 50%) at 150 mM and 300 K. (G)-(I) Intermolecular interaction summary for (G) TDP43-U40 interactions in (50%, 50%), (H) TDP43-GP60 interactions in (50%, 50%), and (I) TDP43-PA60 interactions in (50%, 50%) at 150 mM and 300 K. The fraction of interactions, $\mathsf{F}_{\mathsf{int}}$, are aggregated by type and normalised by the total number of the intermolecular interactions in (A)-(C) respectively.

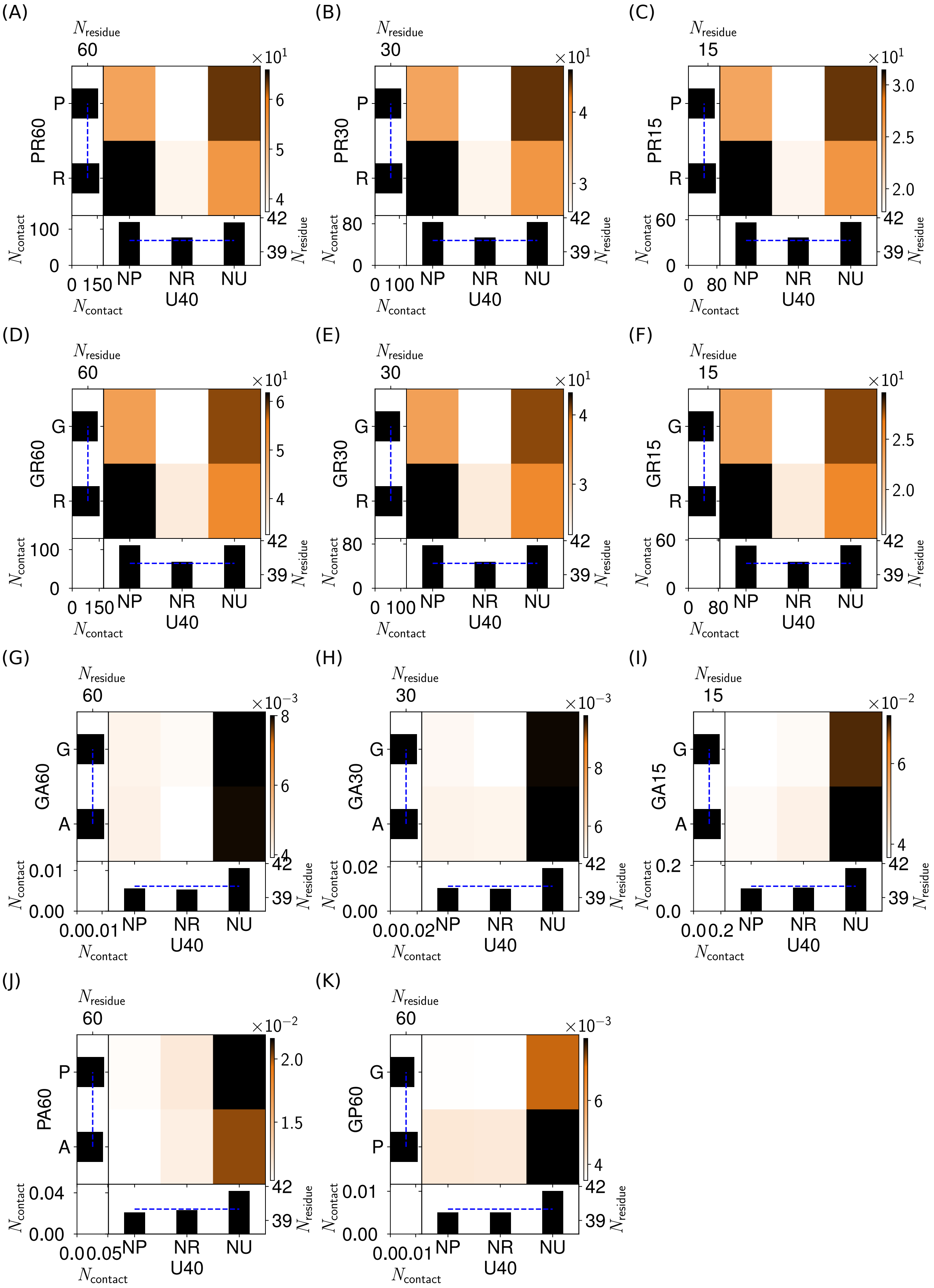


Figure S14: Intermolecular interaction summary for A) U40 with PR60 (50%, 50%) B) U40 with PR30 (50%, 50%) C) U40 with PR15 (50%, 50%) D) U40 with GR60 (50%, 50%) E) U40 with GR30 (50%, 50%) F) U40 with GR15 (50%, 50%) G) U40 with GA60 (50%, 50%) H) U40 with GA30 (50%, 50%) I) U40 with GA15 (50%, 50%) J) U40 with PA60 (50%, 50%) K) U40 with GP60 (50%, 50%) at 150 mM and 300 K.

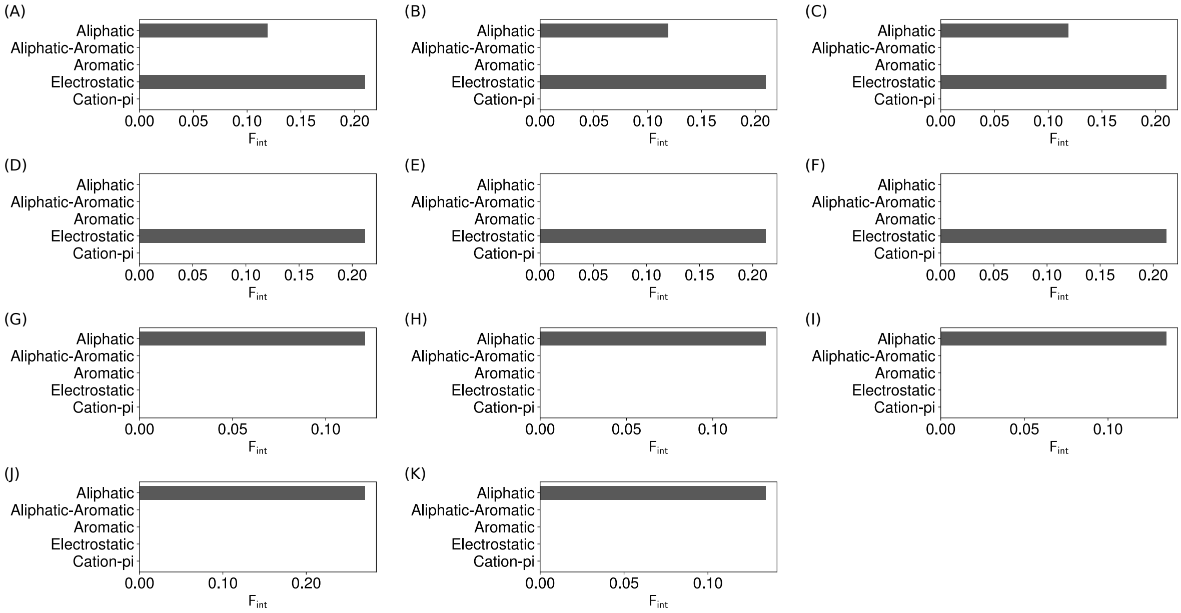


Figure S15: Intermolecular contact map by residue type for A) U40 with PR60 (50%, 50%) B) U40 with PR30 (50%, 50%) C) U40 with PR15 (50%, 50%) D) U40 with GR60 (50%, 50%) E) U40 with GR30 (50%, 50%) F) U40 with GR15 (50%, 50%) G) U40 with GA60 (50%, 50%) H) U40 with GA30 (50%, 50%) I) U40 with GA15 (50%, 50%) J) U40 with PA60 (50%, 50%) K) U40 with GP60 (50%, 50%) at 150 mM and 300 K.
